# Validation of a novel immersive Virtual Reality setup with responses of wild-caught freely-moving coral reef fish

**DOI:** 10.1101/2022.11.01.514726

**Authors:** Manuel Vidal, Suzanne C. Mills, Emma Gairin, Frédéric Bertucci, David Lecchini

## Abstract

Virtual Reality (VR) enables standardised stimuli to invoke behavioural responses in animals, however, in fish studies VR has been limited to either basic virtual stimulation projected below the bowl for freely-swimming individuals or a simple virtual arena rendered over a large field-of-view for head-restrained individuals. We developed a novel immersive VR setup with real-time rendering of animated 3D scenarios, validated in a proof of concept study on the behaviour of coral reef post-larval fish. Fish use a variety of cues to select a habitat during the recruitment stage, and to recognize conspecifics and predators, but which visual cues are used remains unknown. We measured behavioural responses of groups of five convict surgeonfish (*Acanthurus triostegus*) to simulations of habitats, static or moving shoals of conspecifics, predators, and non-aggressive heterospecifics. Post-larval fish were consistently attracted to virtual corals and conspecifics presented statically, but repulsed by their predators (bluefin jacks, *Caranx melampygus*). When simulated shoals passed nearby repeatedly, they were again attracted by conspecifics showing a tendency to follow the shoal, whereas they moved repeatedly to the back of the passing predator shoal. They also discriminated between species of similar sizes: they were attracted more to conspecifics than butterflyfish (*Forcipiger longirostris*), and repulsed more by predators than parrotfish (*Scarus psittacus*). The quality of visual simulations was high enough to identify between visual cues – size, body shape, colour pattern – used by post-larval fish in species recognition. Despite a tracking technology limited to fish 2D positions in the aquarium, preventing the real-time updating of the rendered viewpoint, we could show that VR and modern tracking technologies offer new possibilities to investigate fish behaviour through the quantitative analysis of their physical reactions to highly-controlled scenarios.

## Introduction

Animal behaviours such as foraging, habitat choice, predator avoidance, social behaviour and mate choice, are studied for multiple reasons including to understand how they have been shaped by natural selection or how they are impacted by internal and external stimuli. Animal behavioural studies face two challenges: one is related to the subject’s own understanding of the task to be performed, and the other is linked to measuring its response (Drew, 2019). One way to solve the first challenge is to use a task that requires a natural reaction to a stimulation presented in an ecological context, and for which the understanding is implicit. Conventional experimental approaches used live stimulus animals or environments, but they suffered from a lack of control and standardization as neither the behaviour of stimulus animals nor the local environment can be completely controlled. For instance, testing behavioural dominance in response to an opponent requires trials with multiple opponents of known dominance and applying a correction factor (Alatalo et al., 1991; Mills et al., 2007). Furthermore, experiments with live stimulus animals often require long methodological preparation, which limits the number of possible manipulations (Neri, 2012). As a result, stimuli have been artificially designed to provide repeatable behavioural observations (Carmichael, 1952) and have evolved from simple pictures and physical models to Virtual Reality (VR) that enables standardised manipulations of stimulus behaviours or environments. The second challenge, response measurement, has been solved using video-based tracking systems of freely moving focal animals, which assumes that the behavioural response lies in the kinematics of the animal, e.g. position, orientation, speed, spatial dispersion. Therefore VR provides a good methodological compromise between a perfectly controlled but not ecologically valid stimulation, and a realistic natural situation with little to no parameter control. Although VR simulators have been widely used over the last 25 years to elucidate the perceptual, sensorimotor and cognitive mechanisms underlying human spatial orientation in the environment (e.g. Tarr & Warren, 2002; Vidal et al., 2004; Mossio et al., 2008; Vidal et al., 2009; Vidal & Bülthoff, 2009), only recently have they been adapted to investigate animal behaviour, ranging from mice to fruit flies and zebrafish (Harvey et al., 2009; Stowers et al., 2017; for a discussion, see Drew, 2019).

The first studies of fish visual behaviour that used pre-recorded video stimuli in mating preference tasks date from the end of the nineties, with either manipulated real-videos for (Rosenthal & Evans, 1998) or synthetically generated videos of 3D animated fish (Künzler & Bakker, 1998). Ten years later, the same team showed that computer animations of artificial fish allow manipulating movement, body shape and skin-colour to investigate preferences in the cichlid *Pelvicachromis taeniatus* (Baldauf et al., 2009). The survival potential of prey group formation and movement was measured through the response of real predatory bluegill sunfish (*Lepomis macrochirus*) to virtual prey projections (Ioannou et al., 2012). Both experiments used one or two screen monitors to display the virtual images. Since then, technology has greatly improved, and VR has led to considerable advances in the understanding of the neural bases of zebrafish visual behaviour (Portugues & Engert, 2009; Dunn et al., 2016), shoaling behaviour and social interactions (Larsch & Baier, 2018; Huang et al., 2020; Harpaz et al., 2021), as well as decision making (Barker & Baier, 2015). However, to date the use of VR to study fish behaviour has been restricted to zebrafish larvae, either moving freely in a bowl responding to basic virtual stimulation projected below such as moving dark disks, a checker board or grass bottom, coupled with infrared 3D tracking (Stowers et al., 2017), or with head-restrained zebrafish responding to conspecifics in a simple virtual arena covering 180° of the visual field and rotating based on tail movements (Huang et al., 2020). Therefore, modern VR technology including realistic rendering and immersion in a large 3D volume has not been adapted to fish studies yet, despite the limitless number of findings that can be generated in terms of quantitative animal behaviour and their ecological implications. Here, we carry out a proof of concept study on a new setup for freely-moving fish within an aquarium with an immersive full-field rendering of virtual scenes using projections not only from below, but on all five sides (except the top). We propose that our VR setup has considerable future potential for all types of behavioural studies on fish species at any stage of their life-cycle. Here, our methodology is tested on the behaviour of post-larval coral reef fish exposed to multiple scenarios during their recruitment.

In all marine environments, one of the main mysteries of fish ecology is how larvae recruit onto the relatively rare patches of coastal habitats (for review, see Doherty, 2002; Barth et al., 2015). The life-cycle of most reef fish species starts with a planktonic larval phase, lasting several weeks, followed by recruitment and a sedentary reef phase for juveniles and adults (Leis & McCormick, 2002). At the end of the pelagic phase, this recruitment relies on the detection of a suitable habitat which will facilitate larval survival and growth (Doherty, 2002; Lecchini & Galzin, 2003). Simultaneous to that choice, species-specific changes in morphology and physiology, metamorphosis, occur. These changes are linked to ecological shifts with modifications of diet and diel activity period (McCormick et al., 2002; Besson et al., 2017; Holzer et al., 2017) but also of the sensory systems (Lecchini et al., 2005; Tettamanti et al., 2019). Many studies have highlighted the role of sensory and swimming mechanisms in larval habitat selection, most focusing on the role of chemical (e.g. Atema et al., 2002; Vail & McCormick, 2011; Coppock et al., 2013; Lecchini et al., 2013) and acoustic cues (e.g. Tolimieri et al., 2004; Montgomery et al., 2006; Holles et al., 2013; Parmentier et al., 2015). However, vision is a well-developed sense in coral reef fish larvae (Myrberg & Fuiman, 2002), effective to up to 10 m for *Plectopomus leopardus* post-larval fish at recruitment (Leis & Carson-Ewart, 1999). Once larvae are close to a reef, visual cues of conspecifics become important in the recruitment process (Booth, 1992; Barth et al., 2015). However, only a few studies have identified the visual parameters used by larvae to recognize conspecifics or predators (e.g. Leis & Carson-Ewart, 1999; Booth, 1992; Huijbers et al., 2012; Lecchini et al., 2014). To test how post-larval fish (i.e., larvae having recruited onto a habitat, with metamorphosis still on-going; see Besson et al., 2020) interpret a range of sensory cues, behavioural experiments can reproduce and control a large variety of combinations of visual cues (Barth et al., 2015). VR is potentially an excellent method to test such behaviours as visual factors such as size, colour patterns, and the behaviour of other individuals can be tightly controlled (Stowers et al., 2017; Brookes et al., 2020). Here, we experimentally validate a new and fully immersive VR setup for fish by testing several presentation scenarios, named trials, in three experiments on post-larval fish during recruitment.

We used an innovative immersive VR setup to understand how post-larval convict surgeonfish (*Acanthurus triostegus*) visually recognize a suitable habitat, adult conspecifics and one of their predators (bluefin jacks, *Caranx melampygus*). Our first objective was to experimentally validate the use of simulated 3D models of fishes in a VR setup by confirming that they are realistic enough to cause natural reactions in post-larval fish. Three main experiments were carried out to identify the visual cues used by *A. triostegus* post-larvae to recognize adult conspecifics and a predator. Trials included the presentation of virtual habitats and virtual monospecific fish shoals of conspecifics or predators. The trials with virtual fish species were projected either static (moving in place, Experiments 1 and 2) or dynamic (swimming past on one side of the aquarium, Experiments 2 and 3) and a coral reef habitat on all other sides, aimed to virtually reproduce previous studies in which the reaction to either static fish in the corners of the aquarium (Katzir, 1981) or to real fish swimming in a separate adjacent aquarium (Roux et al., 2016) had been studied. We also tested post-larval behavioural responses to two virtual fish shoals swimming past on either side of the aquarium, each with different species, inducing a forced choice (Experiment 3). Furthermore, this experiment was designed to test whether post-larvae can discriminate between the size and species of virtual fish, *i.e.,* if post-larval fish consider larger virtual fish as threats irrespective of species. The advantage of our virtual presentation compared to using live stimuli, is that we were able to measure post-larval fish behaviour rapidly in response to different trials. Our second objective was to validate the automation of post-larval fish position tracking within the test aquarium at high temporal resolution, enabling detailed characterization of their behavioural responses to each scenario.

### The Immersive VR Setup

The experimental setup was composed of three connected modules: the focal aquarium in which post-larvae could swim freely suspended inside the test aquarium, the rendering module which projected the interactive 3D virtual environments depicting a subaquatic natural scene with fishes and corals, and the tracking module which recorded post-larval behaviour in real-time. The software was developed in the lab and the hardware was assembled by Immersion^TM^.

### Test and focal aquaria

The test aquarium was a rectangular prism made of 10 mm-thick Plexiglas plates, with a 50×50-cm square bottom and 35 cm-high lateral sides. The external faces of the bottom and lateral sides were covered by a retro projection translucent, but not transparent, film, as such post-larval fish inside the aquarium could not see the room surrounding the setup, except for the ceiling. The aquarium was filled with 78 litres of sea water so that the water surface was aligned with the upper limit of the video projection. The entire setup was mounted on a structure made of 4-cm squared-section aluminium bars (**Figure 1A**). A smaller focal aquarium (dimensions 20×20×20 cm) in which the post-larvae were placed, was attached to the structure using chains and positioned inside the test aquarium (**Figure 1B**). This smaller focal aquarium limited post-larval movement maintaining them within the range where geometrical projection distortion and image corrections were minimal and would not affect post-larval behaviour (see Video-based tracking section).

**Figure 1.**
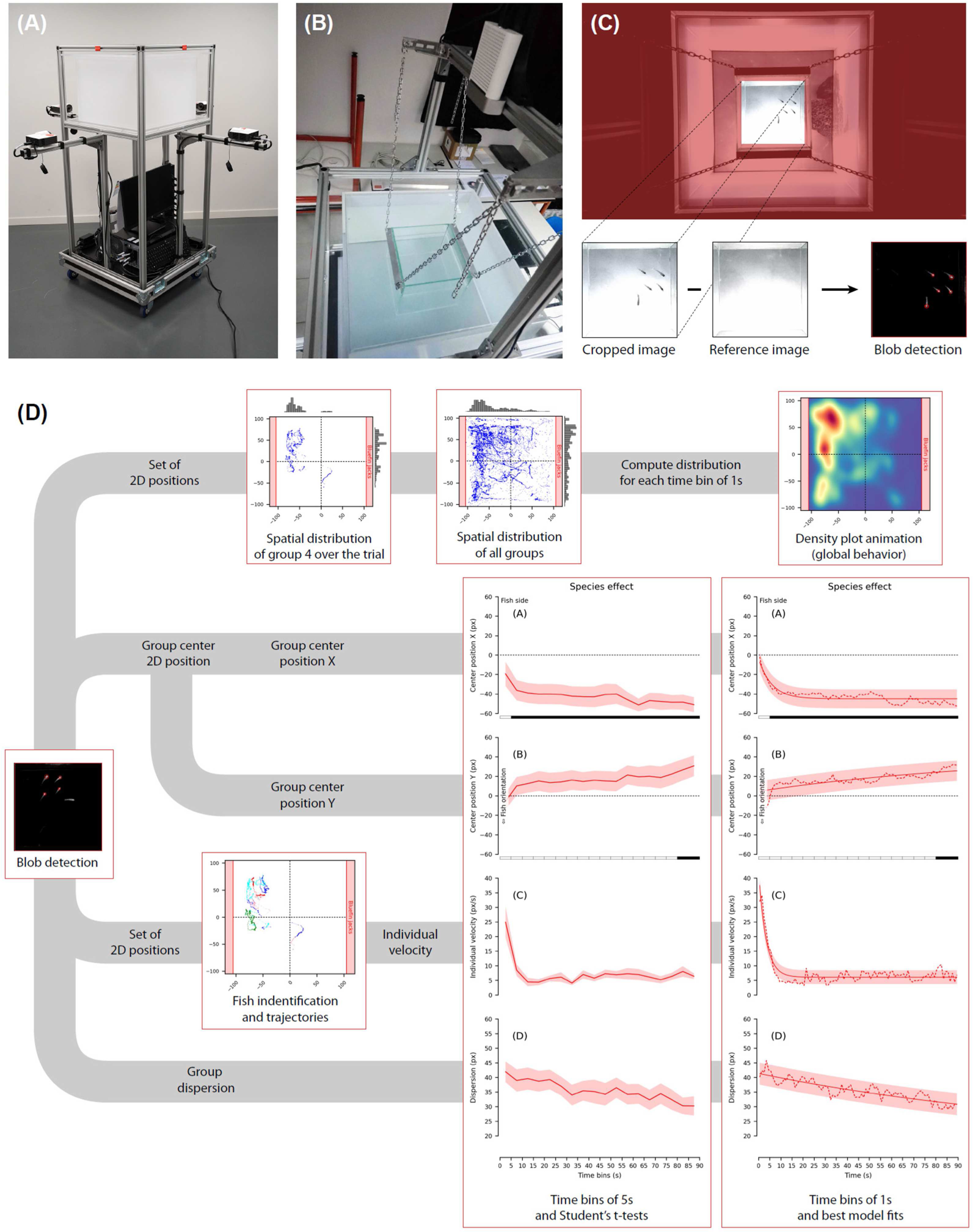
**(A)** View of the experimental setup as delivered by Immersion^TM^ with the test aquarium (dimensions 50×50×35 cm) and 5 video projectors (four lateral sides plus bottom). **(B)** View of the smaller focal aquarium (20×20×20 cm) placed inside the test aquarium in order to limit the displacement range of fish, and of the Microsoft^TM^ Kinect Azure camera recording the behavioural responses. **(C)** Illustration of the detection pipeline starting from the camera view to blob detection executed after cropping and subtracting the reference image, followed by the tracking post-processing pipeline **(D).** Detected 2D positions were used to characterize post-larvae behavioural responses. For each condition, the overall behaviour obtained combining data from all tested groups is visible in the animation of the 2D density heat maps generated every second. The X and Y positions of the centre of the groups, the individual velocities and the dispersions were averaged across time-bins of 5s to perform statistical comparisons (Student’s t-tests), and across time-bins of 1s to fit the behavioural models (regressions). Individual velocities are extracted from the reconstruction of the trajectories, which was based on the identification of each fish from one frame to another (see text).

### Virtual scene rendering

Five video projectors ensured an immersive full-field rendering of the virtual scenes on five sides of the test aquarium (Optoma ML1050ST+, running at 60Hz with a resolution of 1280×800 for the side views and 800×800 for the bottom view). The visible range of post-larval *A. triostegus* likely falls within the human visible range, enabling the use of these video projectors for visual stimulation (Losey et al., 2003). The baseline 3D virtual environment, which was projected on all five aquarium sides at all times, consisted of a sandy bottom at 2 m, with simulated surface ripples and caustics projected on the ground. The virtual viewpoint (position of rendering cameras), which defines the physical-to-virtual relationship, was placed at a depth of 0.75 cm and corresponded to the centre of the test aquarium. Different scenarios (trials) consisting of coral pinnacles (healthy or bleached) as well as animated fish of various species were added to the baseline 3D environment, depending on the simulation, on the left or right side of the aquarium, at a distance of 50 cm. Simulated shoals of five fish could either swim in place on either side or follow a tangent trajectory at a given speed. We used Epic Games Unreal^TM^ Engine 4.23 to render the virtual fish and scene according to the desired test conditions, and to manage the sequencing of trial executions. **Figure 2A** shows how three live post-larvae view a virtual scene of corals and adult surgeonfish from within their focal aquarium.

**Figure 2.**
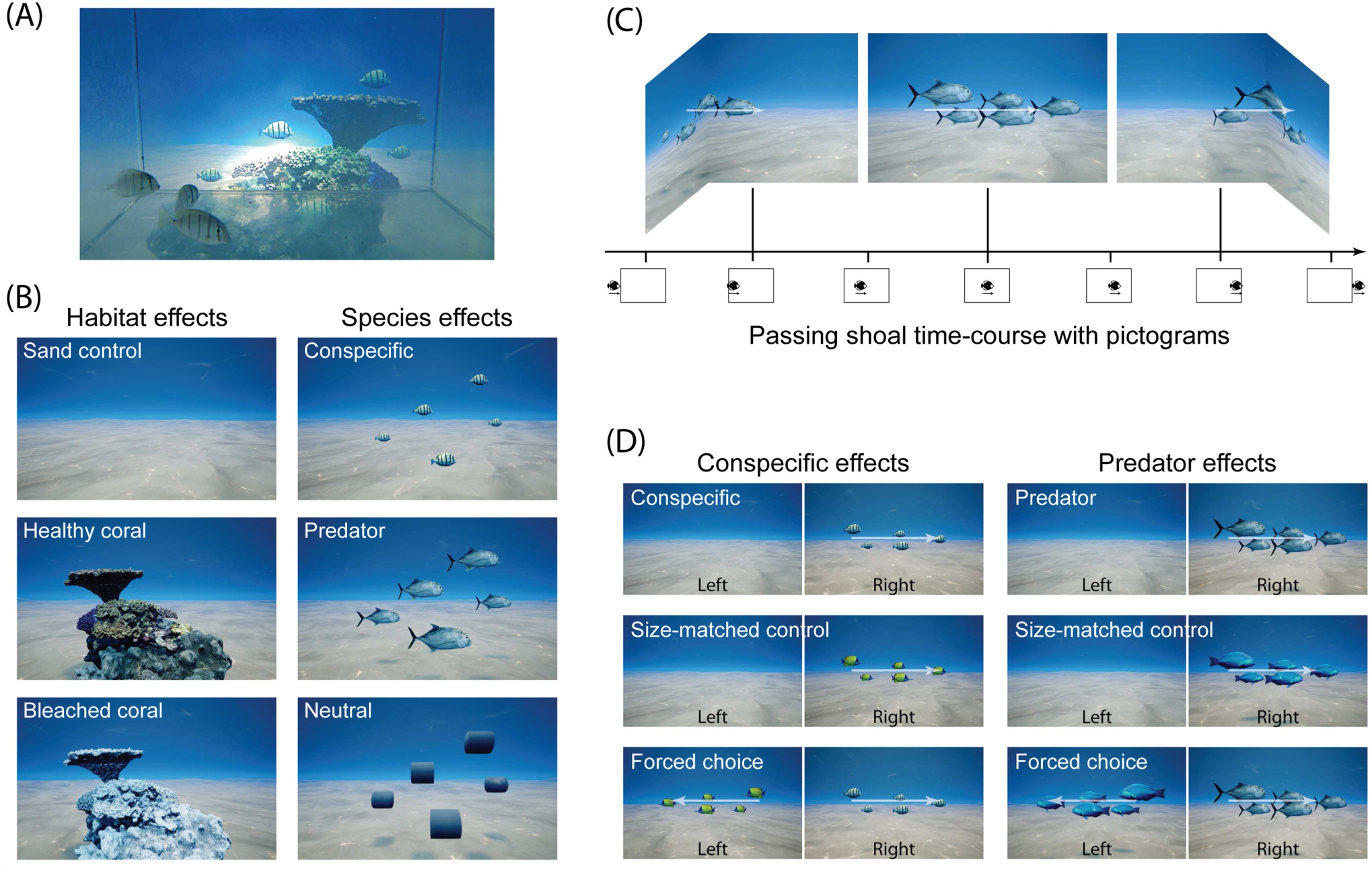
Illustration of the stimuli used in the experiments. **(A)** Rendered scene with virtual corals and adult surgeonfish as seen from inside the focal aquarium by real larvae. **(B)** Experiment 1. Habitat effects (left): Sand control, Healthy or Bleached coral (pinnacles). Species effects (right): Conspecific (adult surgeonfish, *Acanthurus triostegus*), Predator (bluefin jacks, *Caranx melampygus*), Neutral (untextured cylinders). The fish stimuli were presented in shoals of five individuals swimming in place, centred at a virtual location corresponding to 25 cm behind one of the sides of the test aquarium. **(C)** Dynamic presentation of a virtual fish shoal swimming past the post-larval fish (either 1 pass or 3 passes for Experiment 2 and only 1 pass for Experiment 3). The virtual fish species were either conspecifics or predators (as shown here). Shoals swam along a 4.5-m straight line for 30 seconds at 15 cm/s (slow pace). **(D)** Experiment 3. Three conditions were tested to examine the response of the post-larval fish to conspecifics (left): Conspecific alone, Conspecific-sized control alone (butterflyfish, *Forcipiger longirostris*), and Forced choice of conspecific *vs.* conspecific-sized control. Three conditions were also tested to examine the response of the post-larval fish to predators (right): Predator alone, predator-sized Control alone (parrotfish, *Scarus psittacus*), and Forced choice of predator *vs.* predator-sized control. Similarly to Experiment 2, the virtual shoals swam along a 6-m straight line for 40 seconds at 15 cm/s (slow pace).

### Video-based tracking

A Microsoft^TM^ Kinect Azure depth camera was placed 50 cm above the water surface to continuously monitor post-larval fish behaviour in the focal aquarium (**Figure 1B**). For each trial, a top-view colour video was recorded and processed in real-time at a frequency of 5Hz to compute the 2D location of each post-larval fish. In the original design of the setup, we planned to track real-time 3D positions of fish with the infra-red (IR) sensor of the Microsoft Azure Kinect depth camera. However, this technology revealed not suitable for underwater tracking due to the large hot-spot created on the surface of the water by the IR grid-spot. For this reason, we switched to the color sensor and could not adjust in real-time the rendering viewpoint according to fish position in the aquarium. The detection pipeline used both the OpenCV image processing library and the Kinect Azure SDK (**Figure 1C**). Images were extracted from the colour video stream and cropped. The constant background was then removed by subtracting a reference image captured before placing fish in the aquarium. The 2D location of each fish was detected using the OpenCV blob detector with parameters adjusted appropriately. The computer performing the virtual rendering also executed this processing pipeline in real-time. The processing had no impact on the frame rate of the visual scene. Tracking performance is provided in Appendix **Fig A1**.

### Tracking post-processing

The 2D positions of each post-larval fish in groups of 5 were detected at a sampling rate of 5Hz. This automated process is not error-free: in some frames, fewer than five blobs were detected (low signal for smaller juveniles swimming at greater depth or due to overlaps) or more than five blobs (fish reflecting on the Plexiglas when swimming close to the aquarium sides). The tracking post-processing pipeline using Python scripts involved seven steps (**Figure 1D**). First, frames for which only one or two fish were detected were removed to reduce group mean value noise. Second, reflection biases mentioned above were limited by removing the outermost blob when a pair of fish and wall-reflected fish was potentially detected (*i.e.,* when two vertically-or horizontally-aligned blobs were very close to each other and to the edge of the aquarium). Third, as the automated detection cannot identify and track individual fish from one frame to the next (identification problem), a minimal heuristic distance was used to track the fish. This distance was only used when the position change of a blob from one frame to the next was minimal (with five individuals, there were 120 possible combinations across each pair of frames). This provided the (partial) trajectories and instantaneous velocities of each individual. Transiently missing or extra blobs could produce artificial jumps to distant locations; above a given distance threshold between frames (corresponding to 15cm/s) these jumps were ignored in the computation of individual instantaneous velocities. The trajectory reconstructions are plotted with coloured lines for each tracked fish in a (X, Y) square graph representing the aquarium. Fourth, for each validated frame, the X and Y position of the centre of the group, the dispersion relative to the centre (mean distance to the centre), and average individual velocities were computed. To account for the fact that the stimulations were presented either on the right or the left, the sign of the X coordinates was inverted when the stimulation was presented on the left. Fifth, in order to visualize the raw results for each experiment and each tested condition, 2D scatterplots with all valid fish positions from all groups, and normalized X- and Y-position distribution histograms (with 5-pixel large bins) are used (representative scatterplot from Experiment 1 is in **Fig A3**). Sixth, in order to visualize the average behavioural responses across time for each experiment and tested condition, heatmaps of fish position density in the aquarium were plotted in successive 1-second intervals. Lastly, the time-series for each of the four behavioural measures (group centre X- and Y-position, dispersion and individual velocities) were binned into 5-second intervals for the statistical tests, and into 1-second intervals to find the best behavioural model fit.

### Experimental Validation

Three main experiments were carried out to understand how post-larval convict surgeonfish (*Acanthurus triostegus*) visually recognize a suitable habitat, adult conspecifics and one of their predators (bluefin jacks, *Caranx melampygus*). Our first objective was to experimentally validate the use of simulated 3D models of fishes in a VR setup by confirming that they are realistic enough to cause natural reactions in post-larval fish.

## Methods

### Specimen collection

Over 200 post-larval *Acanthurus triostegus* (TL = 2.55-2.75 cm) were captured using hand nets at night, shortly after entering the north-eastern reef crest of Moorea, French Polynesia (17°29′52.19″S, 149°45′13.55″W). Individual *A. triostegus* had not yet acquired skin stripes which only form after recruitment, therefore they were still undergoing metamorphosis, and were considered ‘post-larvae’ (Besson et al., 2020).

### Ethical note

Ethical approval for the study was granted from The Animal Ethics Committee, Centre National de la Recherche Scientifique (permit number 006725). This study also complies with the rules defined by the Direction de l’Environnement de la Polynésie Française (DIREN) regarding experiments on coral fish in aquaria. After captured, post-larval fish were placed in acclimatization aquaria at CRIOBE for 36 hours, in groups of 40 maximum, filled with UV-sterilized and filtered (10-μm filter) seawater maintained at 28.5 °C, under a 12:12 LD cycle. Stress was minimized during transport using occluded small aquaria. Once the experiment was over, animals were returned to their natural habitat.

### Experimental protocol

The behavioural response to the multiple trials – different habitats or fish shoals – was assessed for groups of five post-larval fish placed together in the aquarium. A neutral, baseline 3D environment (sandy bottom with animated caustics) was displayed throughout the experimental sessions on all five sides of the aquarium. Virtual fish or coral pinnacles appeared and disappeared at specific times and in specific virtual locations depending on the trial. Each experiment started with a 4 min habituation period in the baseline environment, followed by trials each lasting 90 seconds (experiment 1 and 2) or 60 seconds (experiment 3). To minimize interference, the baseline environment was also displayed for 2 min between trials. For each experiment, the presentation order of trials was randomised and balanced to avoid order effects and allow for statistical comparisons between pairs of conditions. To exclude possible side biases from the random sand texture pattern or from the room’s ceiling and lightning, the stimulation side was randomly balanced between the left and right of the aquarium. Lastly, after every half-day the aquarium was emptied, washed with freshwater, and refilled and the focal aquarium was oxygenated between replicates.

#### Experiment 1. Effects of static presentation of habitat and fish

Groups of five post-larval fish were presented with six trials: three virtual habitats and three virtual fish or neutral shapes on only one side of the aquarium (randomised) each for 90 seconds (**Figure 2B**). Virtual simulations rendered the post-larval fish at a depth of 1.25 m. The three virtual habitats tested were: Sand control (only the sandy baseline environment); Healthy coral (a pinnacle with healthy tabular and branched corals); and Bleached coral (the same pinnacle, but all corals were bleached). The three virtual fish species consisted of shoals of five virtual fish swimming in place in the same pattern and position: Conspecifics (five adult convict surgeonfish, *Acanthurus triostegus*); Predators (five bluefin jacks, *Caranx melampygus*); and Neutral (five large untextured cylinders). The individual positions of five post-larval fish were constantly tracked during the six successive trials presented in the following order: Sand control (1^st^); Healthy or Bleached coral (randomly 2^nd^ or 3^rd^); Conspecific, Predator or Neutral (randomly 4^th^, 5^th^, or 6^th^). Sixty post-larval fish were tested in 12 groups of 5 fish, and for each the total experimental duration was approximately 22 minutes.

#### Experiment 2. Effects of static versus dynamic presentation of fish

In Experiment 2, with 8 trials, behavioural responses to the static presentation of shoals of five virtual fish was compared with behavioural responses to a more realistic dynamic situation in which shoals of five fish appeared, swam past the post-larval fish in a non-aggressive manner and disappeared. Three virtual fish shoal trials were projected: one static shoal swimming in place (as in Experiment 1) for 90 seconds; one shoal swimming by during the first 30 seconds, then disappearing, followed by 60 seconds of the baseline sand environment; or three successive shoals of five fish swimming nearby and disappearing, each over 30 seconds. In each, virtual shoals swam for 30 seconds at 15 cm/s (slow pace) along a virtual line placed 115 cm from the centre of the test aquarium, covering a total distance of 4.5 m (**Figure 2C**). The virtual fish shoals were either surgeonfish conspecifics or bluefin jack predators and were presented in 8 successive trials in the following order: Sand control (1^st^); Conspecific/Predator static, 1 pass, or 3 passes (randomly presented in 2^nd^, 3^rd^, or 4^th^ position); sand control (5^th^); Conspecific/Predator static, 1 pass, or 3 passes (randomly presented in 6^th^, 7^th^, or 8^th^ position). The order in which the fish shoal was presented (conspecifics or predators first) was varied and the side of the aquarium on which the virtual fish were presented was randomised. Sixty post-larval fish were tested in 12 groups of 5 post-larvae, and for each the total experimental duration was approximately 26 minutes.

#### Experiment 3. Effect of size-controlled dynamic presentation of fish

In Experiment 3, behavioural responses to a dynamically swimming shoal of conspecifics or predators, was compared with behavioural responses to size-matched heterospecifics. Six virtual fish shoal trials were projected (**Figure 2D**): surgeonfish conspecifics on one side; conspecific-sized control fish on one side (butterflyfish, *Forcipiger longirostris*); conspecifics and conspecific-sized controls on opposite sides (two-alternative choice); bluefin jack predators on one side; predator-sized control fish on one side (parrotfish, *Scarus psittacus*); predators and predator-sized controls on opposite sides (two-alternative choice). Changes in post-larval fish positions from before to after a virtual shoal swam by were identified. All virtual fish swam at 15 cm/s (slow pace) for 40 seconds along a virtual line placed at 115 cm from the centre of the aquarium (total distance travelled: 6 m). The order in which fish shoal types was presented, conspecifics or predators, was balanced between groups, however, the two-alternative choice was always presented after the single-choice trials. The order of single-choice trials was also balanced. In all conditions, the stimulus was presented over 40 seconds, and post-larval position recording started 10 seconds before and ended 10 seconds after the stimulus (total duration of 60 seconds). Eighty post-larval fish were tested in 16 groups of 5 fish, and for each the total experimental duration was approximately 26 minutes.

### Data analysis

For all experiments, tracking data was sampled at 5Hz, during 90-second trials for Experiment 1 and 2, and 60-second trials for Experiment 3 (see **Figure 1C**). Representative trajectories of post-larval groups are available in Appendix **Fig A2**, **Fig A7** and **Fig A11** for Experiment 1, 2 and 3, respectively. Animated heatmaps of fish position density in the aquarium for all conditions are available in Appendix **Video A4**, **Video A8** and **Video A12** for Experiment 1, 2 and 3, respectively. Note that for all 2D plots, data is organized so that the simulation is always presented on the right side, except for the forced-choice conditions of Experiment 3, for which the stimulation is presented on both left and right sides. General repeated-measures ANOVAs with trial and 5-second time-bin as main factors were generated using the four behavioural measures (group centre X- and Y-position, dispersion and individual velocity). Comparisons between relevant trials (paired Student t-tests) and the deviation from zero of the group’s X and Y positions (Student t-tests against a single value of 0) were conducted for each time bin. The alpha value for significance was adjusted using Bonferroni’s correction for multiple comparisons on a single dataset and visualized in the plots using different grey levels (white for P>0.05 and from light grey for P<0.05/1 to black for P<0.05/n_Tests_ with n_Tests_ = 3 for either the 3 paired comparisons or the 3 single-value comparisons). Plots displaying the timeseries of the four behavioural measures (binned in 5-second intervals) and the results from the statistical comparisons are provided in the Appendix **Fig A5**, **Fig A9** and **Fig A13** for Experiment 1, 2 and 3, respectively.

### Behavioural model fit

To characterize the temporal aspect of post-larval fish behavioural responses, we designed several models taking into account the distance of the post-larval fish to the simulated shoals. The position of the virtual simulations was sustained and constant throughout the trials in Experiment 1 and in the static trials of Experiment 2, but the virtual shoals moved in the dynamic conditions of Experiments 2 and 3. Because of this major difference between the static and dynamic conditions, we used a different set of possible behavioural models for either type of trial to measure behaviour (centre position, individual velocity and dispersion). For the static conditions, we tested three simple ecologically relevant models (linear, quadratic and exponential) and for dynamic conditions, we added a periodic component to capture the cyclic variations of the stimulation (**Table 1**). The average behavioural measures obtained for the tested groups (n=12 or n=16), binned in 1-second intervals, were fitted using each of the three models. The fit quality was estimated with the root mean square error (RMSE) between the average data points and the model predictions. In order to avoid data over-fitting, we used a limited number of models and selected the best model based on the trade-off between fit quality (RMSE) and the number of parameters. Linear models (for static conditions) and linear periodic models (for dynamic conditions) have one parameter fewer than the quadratic and exponential models. They were favoured when the RMSE difference with the other models was below 5% (*e.g.,* if the linear and quadratic fits had RMSEs of 10 and 9.6 resp., the linear model was selected). Lastly, in order to check the validity of each fit, the obtained RMSE was compared to the RMSE distribution obtained by applying, for the given condition and measure, the same fitting procedure but with scrambled time-bins a thousand times. Two quality criteria were used: if the obtained RMSE was lower than 0.8 times the mean RMSE distribution, and below the lower 1% confidence interval bound of the distribution, the fit was considered valid (good signal-to-ratio level). The results are summarized in plots displaying, for each condition, the time-series of each measure in bins of 1 second, the best behavioural model fit, and the outcome of the statistical comparisons. For each condition, the selected model and adjusted parameters are detailed in Appendix **Table A6**, **Table A10** and **Table A14** for Experiment 1, 2 and 3.

**Table 1.** The sets of behavioural models fitted for the static and dynamic trials. For each model, the equation, the description of the behaviour that it captures, and the meaning of its parameters are detailed.

|  | Name | Equation | Description and meaning of the parameters |
| --- | --- | --- | --- |
| Static conditions | Linear | $a \cdot t + b$ | Linear models capture responses that change constantly in time. These drifts can be observed in slow responses to a potential danger.<br>$a$ : linear term (drift speed)<br>$b$ : offset (start position) |
| | Quadratic | $a \cdot t^2 + b \cdot t + c$ | Quadratic models capture responses that change constantly in time (1<sup>st</sup> order) and that have a reversal (2<sup>nd</sup> order). Reversal are observed of after a loss of interest when attracted by the stimulus ($a < 0$ if in right side) or after habituation to a fearful stimulus ($a > 0$).<br>$a$ : quadratic term (reversal)<br>$b$ : linear term (drift speed)<br>$c$ : offset (start position) |
| | Exponential | $I - (I - s) \cdot \exp(-k \cdot t)$ | Exponential attenuation models capture natural decaying phenomena within a limited spatial or temporal range. These can be observed in quick escape response to a fearful stimulation.<br>$I$ : limit of the attenuation (reached when $t \rightarrow \infty$)<br>$s$ : offset (starting position)<br>$k$ : attenuation factor (positive value, higher for faster decays) |
| Dynamic conditions | Linear periodic | $a \cdot t + b + \frac{1}{2}A \cdot \cos(2\pi \cdot (t - t_0)/T)$ | Linear models combined with a periodic component capture linear drift responses to a cyclic stimulation. Periodic component parameters:<br>$A$ : amplitude of the oscillations (positive value)<br>$t_0$ : phase of the oscillations (peak time, within $[0 \text{ s}, T]$)<br>$T$ : period (**fixed parameter** set to 30 s and 40 s for Experiment 2 and 3) |
| | Quadratic periodic | $a \cdot t^2 + b \cdot t + c + \frac{1}{2}A \cdot \cos(2\pi \cdot (t - t_0)/T)$ | Quadratic models combined with a periodic component capture 1<sup>st</sup> and 2<sup>nd</sup> order responses to a cyclic stimulation. |
| | Exponential periodic | $I - (I - s) \cdot \exp(-k \cdot t) + \frac{1}{2}A \cdot \cos(2\pi \cdot (t - t_0)/T)$ | Exponential attenuation models combined with a periodic component capture naturally decaying periodic responses to a cyclic stimulation. |

## Results

### Experiment 1. Effects of static presentation of habitat and fish

The behavioural responses to the trials were assessed for groups of five post-larval *A. triostegus* placed together in the aquarium. All trials were tested on all groups of post-larval fish. Typical individual trajectories of post-larval fish in response to each of the six trials are shown in Appendix **Fig A2**, scatterplots with all positions occupied by all post-larval fish in Appendix **Fig A3**, and animated heatmaps with the presence density at each successive 1-second intervals in Appendix **Video A4**.

#### Habitat effect

Post-larval fish reactions were similar across habitat types (Sand control, Healthy and Bleached coral). Post-larval fish group centre’s X- and Y-positions were not significantly different between trials either over the whole test period or for most 5-second intervals (lack of significance in boxes below plots in **Figure 3A** and **B**). Apart from a small positive bias (16.5 px) in the Y-position for the Sand control (entire range: t(11)=2.39, P<0.04), possibly related to the initial location in which fish were placed in the aquarium, the X- and Y-positions were not biased to either side of the aquarium (**Figure 3A** and **B**). The level of noise did not allow for good quality fits of linear, quadratic or exponential models (**Table A6**), but the best fit models largely overlapped, confirming non-significant differences in response behaviours across habitat types. In contrast, there was a significant effect of habitat on individual velocities (F_2,22_=5.5, P<0.015, η_G_^2^=0.12), with post-larval fish moving faster (more activity) when presented with Healthy (23.8 px/s, t(11)=2.26, P<0.05) and Bleached (23.7 px/s, t(11)=3.11, P<0.01) corals compared to no corals in the Sand control (14.7 px/s) over the whole time range and during some of the 5-second intervals (shaded boxes below plot in **Figure 3C**). No significant difference in group dispersion – either with statistics or model fitting – was observed across habitat types (**Figure 3D**).

**Figure 3.**
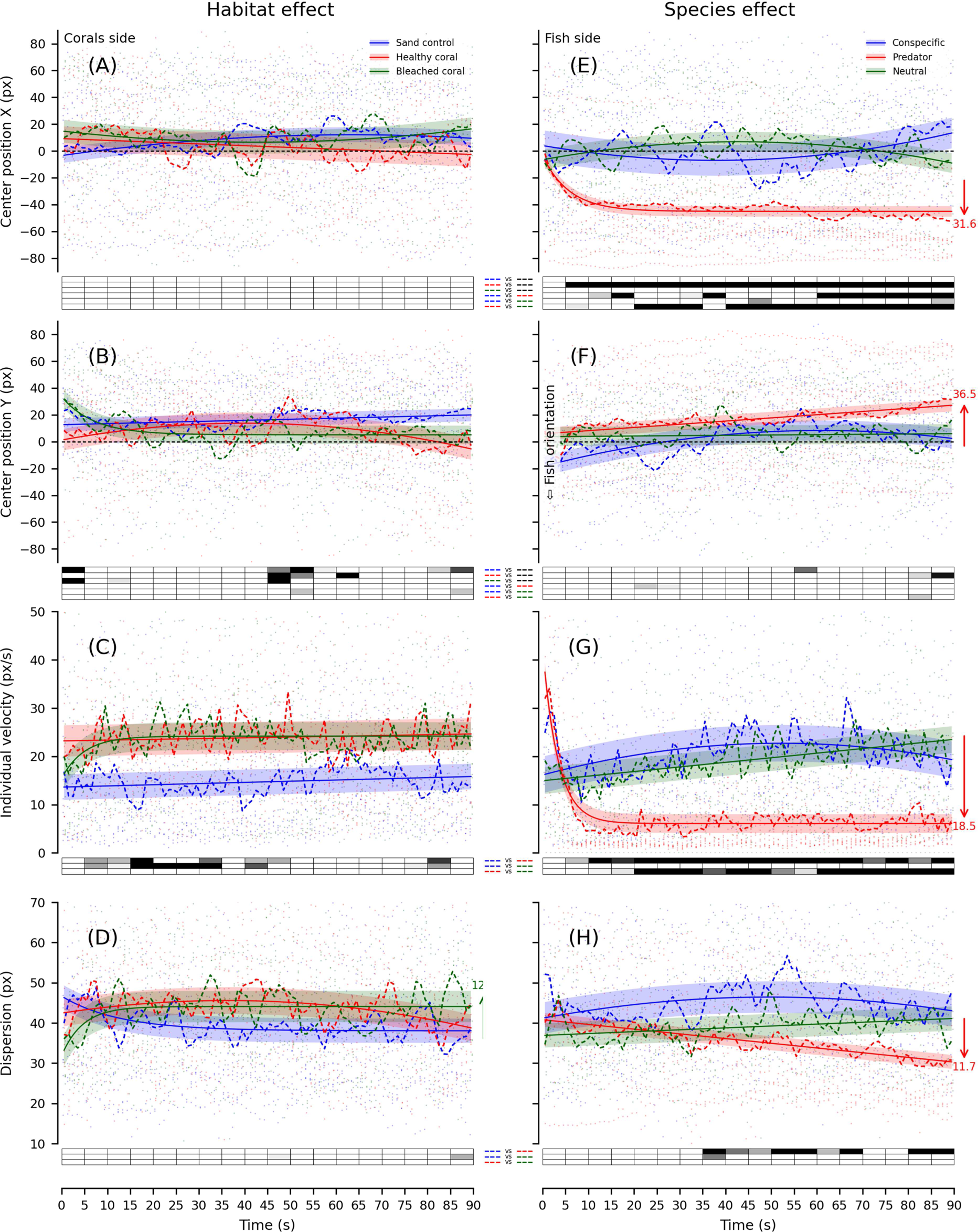
Experiment 1 group time series, with best model fits and statistics. X- and Y-positions of the group centre, individual velocity, and group dispersion for Habitat **(A-D)** and Species trials **(E-H)**. Coloured dashed lines show the average at each time-point of n=12 groups (Habitat trials: Sand control in blue, Healthy coral in red, Bleached coral in green; Species trials: Conspecific in blue, Predator in red, Neutral in green). Coloured lines show the best fitting model for the corresponding trials and error stripes show the RMSE. For each 5-second time-bin, average performances were compared either between each trial or to zero with paired and single-value Student t-tests. Significance levels are provided in the boxes below the plots (ranging from light grey for P<0.05/1 to black for P<0.05/n_Tests_ using Bonferroni’s correction for n_Tests_=3; white for P>0.05).

#### Species effect

Post-larval fish reactions did not differ when presented with static shoals of conspecifics (surgeonfish) and neutral cylinders, but were significantly different when presented with predators (jack fish). Type of fish shoal impacted the post-larval fish group centre’s X-position (main effect, F_2,22_=8.3, P<0.002, η_G_^2^=0.27), which was significantly lower with Predators (–42.3 px) than with Conspecifics (–1.5 px, t(11)=3.14, P<0.01) or Neutral cylinders (1.8 px, t(11)=3.7, P<0.005) across the entire time-range and for most 5-second intervals (shaded boxes below plot in **Figure 3E**). Post-larval fish swam away from the virtual predators: they moved 31.6 px from the first to the last time-bin (t(11)=2.83, P<0.02), mostly at the beginning of the trial (exponential model with *k* parameter of 0.17 s^-1^). On the other hand, with virtual conspecifics post-larval fish hit the sides of the aquarium near the conspecifics more often compared to neutral cylinders or sand control. There was no global effect of the type of virtual species on post-larval fish Y-positions across the entire time-range and for any interval (**Figure 3F**). However, post-larval fish moved slowly toward the upper-left quadrant of the aquarium – the opposite side to the stimulation – and moved to the back of the virtual shoal (linear model with a slope of *a*=0.238 px/s) moving 36.5 px (t(11)=3.09, P<0.01) behind the virtual predators, reducing dispersion (Appendix **Video A4**). Type of fish shoal had a significant effect on individual velocities (F(2,22)=11.08, P<0.001, η_G_^2^=0.20), with lower speeds with Predators (7.3 px/s) than with Conspecifics (20.9 px/s, t(11)=4.09, P<0.002) and Neutral cylinders (19.3 px/s, t(11)=3.23, P<0.01) over the entire time range, and most 5-second intervals (**Figure 3G**). Furthermore, with predators individual velocities rapidly decreased by 18.5 px/s over the first ten seconds (t(11)=3.47, P<0.005), remaining at 6.1 px/s until the end of the trial (exponential model with a very high *k* parameter value of 0.30 s^-1^). Movement, as well as space occupied in the aquarium, were similar when presented with surgeonfish, cylinders or only sand (Appendix **Fig A3**). The effect of fish shoal type on group dispersion was nearly significant (F(2,22)=2.89, P=0.077, η_G_^2^=0.071), due to less dispersion with Predators (35.5 px) than with Conspecifics (44.9 px) across the entire time range (t(11)=3.37, P<0.007), and in most intervals after 35 seconds (**Figure 3H**). With predators, group dispersion decreased slowly and progressively during the trial (linear model with *a*=–0.12 px/s), by 11.7 px (t(11)=2.90, P<0.015): post-larval fish tended to gather after detecting a threat. The best-fitting models of the behavioural reactions to virtual bluefin jacks highlight natural repulsion from a fear-invoking stimulation: the exponential models captured quick responses in a limited space/time range (X-position and individual velocity), whereas the linear models captured slow drifting responses (Y-position and dispersion). In general, the four different behavioural indicators (X and Y position, individual velocity, and dispersion) were not different between conspecifics and neutral cylinders, and despite high variability between individuals resulting in poor quality fits, models also mostly overlapped (**Table A6)**. However, when presented with predators, the quality of model fitting for all behavioural measures was excellent i.e. post-larvae showed homogeneous behaviours within each group as well as across groups.

### Experiment 2. Effects of static versus dynamic presentation of fish

Typical individual trajectories of post-larval *A. triostegus* in response to each of the eight trials are shown in Appendix **Fig A7**, and animated heatmaps with the presence density at each successive 1-second interval in Appendix **Video A8**. Sand control trials were presented before the conspecific and predator trials to provide acclimatization periods. Since no qualitative or statistical differences in behaviour were observed, these results are not reported here.

#### Conspecific effect

Post-larval fish reactions were different when presented with either static (swimming in place) or dynamic (1- or 3-passes) surgeonfish. In the static trial, groups came close to the stimulation (hitting the aquarium side), similarly to Experiment 1, and the group centre’s X-position did not deviate significantly from zero. The best-fitting model quadratic only of average fit but its positive *a* parameter suggested an habituation to an initially slightly fear-invoking stimulus (**Figure 4A** and **Table A10**). In contrast, in the dynamic 1- or 3-pass trials, post-larval fish joined and moved with the virtual surgeonfish shoal as it travelled along the aquarium side (oscillating X- and Y-positions, **Figure 4A** and **B**). When the shoal passed 3 times, post-larval fish kept repeating the same behaviour periodically, without any noticeable attenuation indicating no loss of interest (**Video A8**). The linear periodic model was the best fit, as it captured the post-larval fish cyclic response, with a phase *t_0_* of approximately 17 s for both 1- and 3-pass tests, and an amplitude *A* of 21.6 px (1 pass) and 24.7 px (3 passes). The linear component had an offset *b* of about 14 px for both pass types and a slope *a* of –0.13 px/s (1 pass) and null (3 passes). Oscillations in the X-position showed a significant deviation from zero toward the stimulation side when the shoal was passing (from 15 to 25 s for both 1- and 3-passes, and from 45 to 55 and 75 to 85 s for 3-passes). For the static trial the group centre’s Y-position did not significantly deviate from zero and the best fit was exponential, but of poor quality (**Figure 4B**). For the 1-pass trial, the linear periodic model had the best fit, with a similar phase (*t_0_*=15.6 s) but with a much smaller amplitude (*A*=8.5 px) than for the X-position. The number of post-larval fish that followed the single shoal passage (**Video A8**) was too limited to produce a clear trough, resulting in less vertical motion. Moreover, the linear decay and loss of synchronization in the model after the first cycle was due to the absence of stimulation after 30 seconds. For the 3-pass trial, the best fitting model was exponential and periodic, with a strong attenuation factor of *k*=0.177 s^-1^, rapid oscillations around *l*=7.7 px, a peak at *t_0_*=13.2 s, and large amplitudes (*A*=41.8 px) (**Figure 4B**). The first two oscillations showed significant deviations from zero on peaks and troughs. The X- and Y-oscillations were synchronized and phase-locked with the three virtual surgeonfish shoal passes: vertical motion widely overlapped with the shoal’s linear displacement along the aquarium side, and the horizontal motion corresponded to the movement of post-larvae closer to the shoal at each pass. The swimming kinematic analyses of swimming provide evidence of cohesive group behaviour, in which larvae naturally recognize and follow their conspecifics. Individual velocities were rapidly significantly lower (at 12 px/s then increased linearly, with a fitted slope of 0.064 px/s) with static compared to dynamic virtual conspecifics (**Figure 4C**). This is consistent with an initially slightly fearful response. In dynamic trials, individual velocities oscillated around 20 px/s and decreased to similar values as in static trials towards the end of the trial, but noise was too high for the periodic models to reach a good fit (**Table A10**). Group dispersion tended to be lower for the static than dynamic trials, and fit was poor (**Figure 4D**).

**Figure 4.**
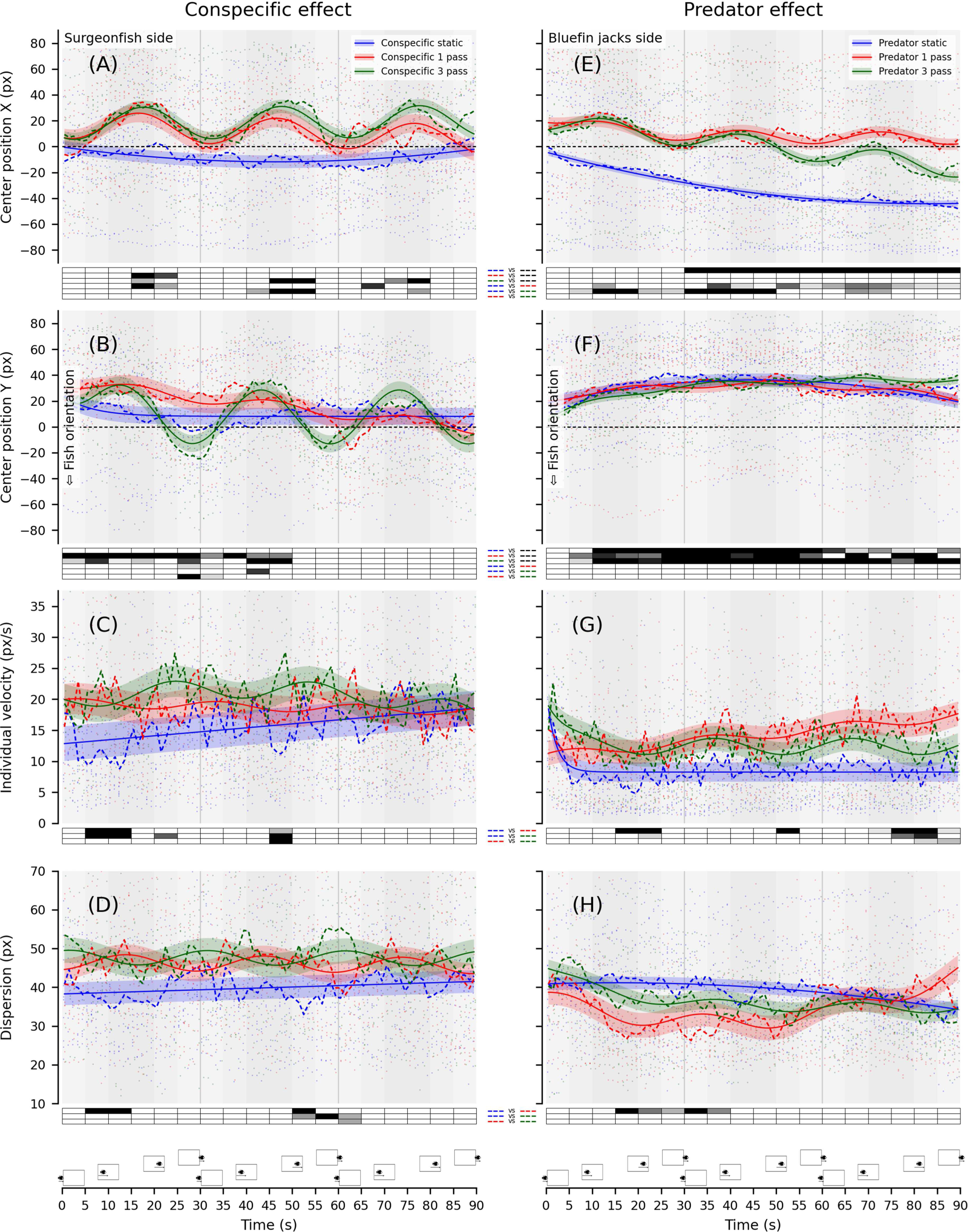
Experiment 2 group time series, with best model fits and statistics. X- and Y-positions of the group centre, individual velocity, and group dispersion for Conspecific **(A-D)** and Predator trials **(E-H)**. Coloured dashed-lines show the average at each time-point of n=12 groups (Conspecifics and Predator trials: static in blue, 1 pass in red, 3 pass in green). Coloured lines show the best fitting model for the corresponding trials and error stripes show the RMSE. For each 5-second time-bin, average performances were compared either between each trial or to zero with paired and single-value Student t-tests. Significance level is provided in the boxes below the plots (ranging from light grey for P<0.05/1 to black for P<0.05/n_Tests_ using Bonferroni’s correction for n_Tests_=3; white for P>0.05). Shaded areas highlight the stimuli critical periods based on the on-going distance of the virtual shoals in the 3-pass conditions (30-second periodicity) illustrated by the fish positions relative to the side screen shown above the time axis.

#### Predator effect

Post-larval fish reacted very differently in response to static compared to dynamic bluefin jack shoals. Static presentation produced the same reaction as in Experiment 1: movement to the opposing side and to the back of the virtual shoal with reduced group dispersion. In the 1- or 3-pass dynamic tests, the density patterns of heatmaps were bimodal, indicative of two reaction types: one similar to that of the static trial, while the second was an asynchronized back-and-forth movement in the X-dimension with the post-larvae moving behind the virtual predators after each pass. The balance between these two reactions varied, with the proportion of static-like behaviour increasing through time. In the static condition, the gradual decrease in X-position (significant deviation from zero in all time-bins beyond 30 seconds) was best fitted with a quadratic model (*a*=0.006 px/s^2^, *b*=– 0.96 px/s and *c*=–4.3 px), with parameters for the climax and reversal outside of the trial time-range (**Figure 4E** and **Table A10**). The best fitting model for the 3-pass condition was linear periodic, with peaks occurring earlier than with conspecifics (*t_0_*=12.8 s), a limited amplitude (*A*=14.6 px), and a linear drift (slope *a*=–0.404) of post-larval fish that gradually moved to the opposing aquarium side. For the 1-pass trial, the response to the first pass overlapped with that of the 3-pass (*t_0_*=12.7 s and *A*=9.5 px) and after this pass, oscillations and drifting were exponentially attenuated. None of the X-positions in the dynamic scenario deviated significantly from zero, but post-larval fish were significantly closer to the stimulation side than in the static trial over multiple time bins. The Y-position was very similar in the three trials (**Figure 4F**): post-larvae moved rapidly behind bluefin jack shoals whether swimming in place (quadratic model) or passing nearby once (quadratic periodic) or three times (exponential periodic). Despite good quality fits, these models largely overlapped and deviated significantly from the midline zero in almost every time bin after the first 5 seconds. At the end of the trials, post-larvae tended to move back towards the midline in the static condition (habituation) and in the 1-pass condition (no more stimulation). Consistent with Experiment 1, individual velocity in the static trial decreased rapidly from 21.3 px/s (*k*=0.50 s^-1^) to 8.2 px/s, which matched an exponential attenuation model (**Figure 4G**). In the 3-pass condition, post-larval velocities fit an exponential periodic model, decreasing rapidly from 19.4 px/s (*k*=0.27 s^-1^) to oscillate around 12.4 px/s at small amplitude (*A*=2.6 px). Furthermore, velocities decreased when post-larval fish noticed predators at each new pass (peak phases locked to *t_0_*=6.5 s). In the 1-pass trial, oscillations had a lower amplitude (*A*=1.5 px), and individual velocities increased linearly after the shoal pass, with underlying oscillations starting once speeds exceeded 10.9 px/s. The only difference between trials occurred towards the end of the static and 1-pass trials, when post-larval fish swam faster in the absence of stimulation. Group dispersion was lower for the dynamic 1-pass than for static trials (15-40 seconds; **Figure 4H**), increasing again in the absence of stimulation (quadratic periodic model). In the static trial, dispersion decreased slowly (quadratic model) but less than in Experiment 1, possibly due to the influence of dynamic trials used in the design. In the 3-pass trial, dispersion was exponentially attenuated, starting at 45 px dropping almost linearly to 34 px (low *k* factor of 0.06 s^-1^).

### Experiment 3. Effect of size-controlled dynamic presentation of fish

Typical individual trajectories of post-larval *A. triostegus* in response to each of the six trials are shown in Appendix **Fig A11**, and animated heatmaps with the presence density at each successive 1-second interval in Appendix **Video A12**. Contrary to previous experiments, model fitting was limited to the interval of 10-50 s, during which the stimulus was visible: the cycle duration of the periodic component was set to 40 s.

#### Conspecific effect

Post-larval fish reactions to virtual conspecific surgeonfish or control size-matched butterflyfish passing nearby were mostly similar, yet with subtle differences. Post-larval fish showed the same behaviour as in Experiment 2: they moved to the upper-right quadrant, but fewer post-larval fish followed the virtual shoal of non-aggressive size-matched controls along the Y-axis compared to conspecifics. Furthermore, dispersion increased faster in the Control compared to Conspecific trials. When both conspecifics and controls were presented (Forced choice), most post-larval fish gathered on the conspecifics side and followed the shoal. The best model for all conditions was linear periodic, except when stated otherwise (**Table A14**). The linear components of the change in X-positions in the Conspecific and Control trials were positive, suggesting an increase in interest over the trial (**Figure 5A**). However, the oscillation during the single pass had a larger amplitude and peaked a few seconds later for surgeonfish (*A*=45.7 px and *t_0_*=31 s) than control butterflyfish (*A*=26.3 px and *t_0_*=24 s). The time interval during which the X-position deviated significantly from the midline was wider for conspecifics (more than three time-bins compared to a single one) and there was a significant difference in X-position between the Control and Conspecific trials during the 35-40 s interval. The linear component of the Y-position in both the Conspecific and Control trials had the same positive slope (*a*≈0.4 px/s) and oscillation phase (*t_0_*≈26 s), but the amplitude in the Conspecific trial was smaller (*A*=26 px against 46 px, **Figure 5B**). The Y-position did not deviate from the midline, and was significantly higher in the Control trial. When both fish shoals were presented (Forced choice), post-larval fish responses were less intense than when presented with only one stimulation but still showed the same two reaction types, *i.e.*, following the conspecifics or remaining in the upper-left quadrant. The X-position was positive and peaked at *t_0_*=38 s, but deviation from midline was not significant due to a strong linear decrease (*a*=-0.92). Altogether, these results indicate that post-larval fish were more attracted to and followed conspecific surgeonfish, whilst spending more time in the upper-right quadrant with similarly-sized butterflyfish. Individual velocity measurements were noisy but remained relatively constant throughout the three trials (at 30 px/s), with fits that mostly overlapped (**Figure 5C**). The linear component of the good quality fit group dispersion model was similar in all three trials, with a nearly null slope (constant) at *b*≈50 px (**Figure 5D**). There were differences in the periodic component, with oscillations of smaller amplitude and troughing 6s later with Conspecifics (*A*=6.6) compared to Controls (*A*=12.2 px), confirming that dispersion increased faster after the butterflyfish shoal passed than after the conspecifics.

**Figure 5.**
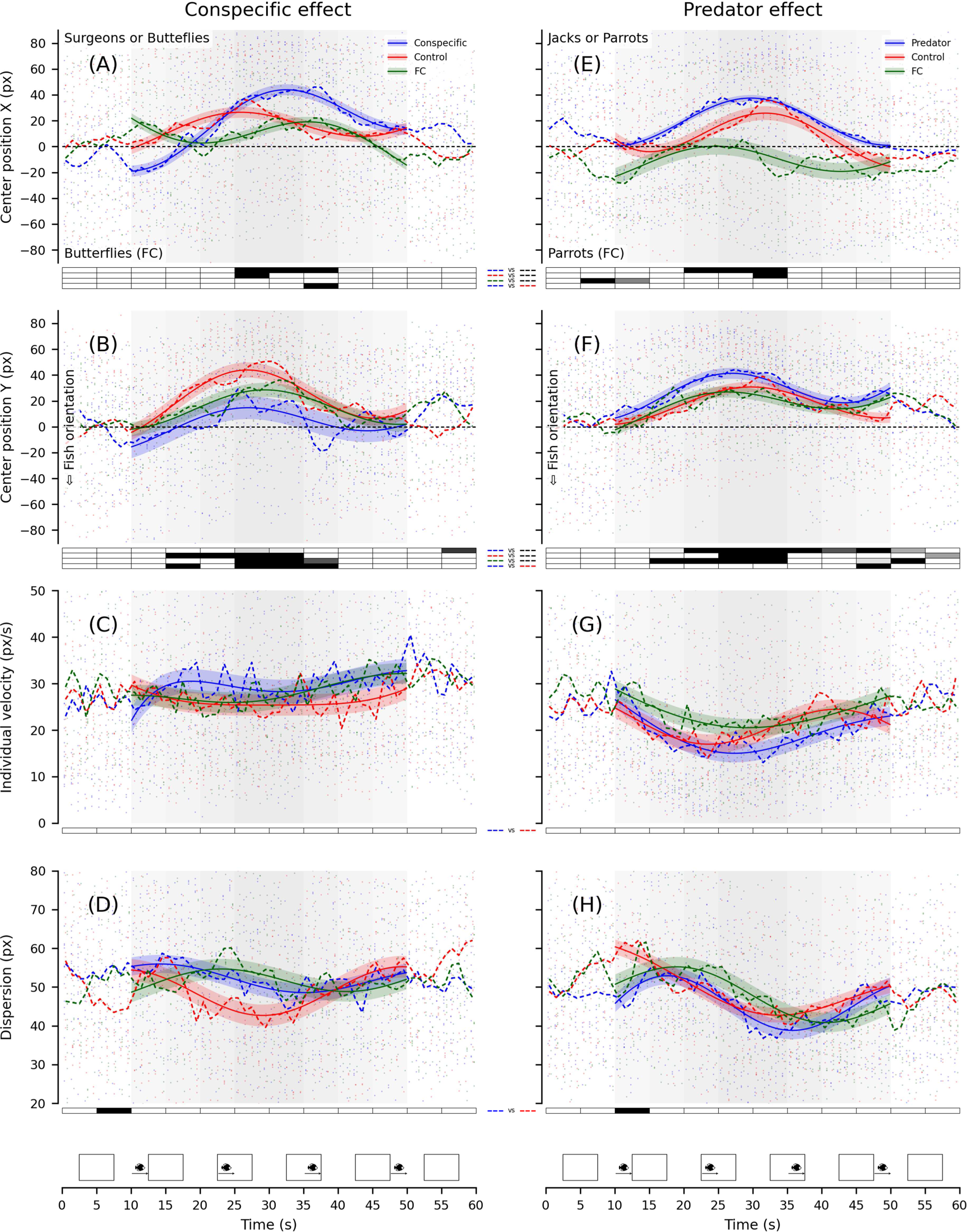
Experiment 3 group time series, with best model fits and statistics. X- and Y-positions of the group centre, individual velocity, and group dispersion for Conspecific **(A-D)** and Predator trials **(E-H)**. Coloured dashed-lines show the average at each time-point of n=16 groups (Conspecific trials: Conspecific in blue, Control in red, Forced-choice in green; Predator trials: Predator in blue, Coloured lines show the best fitting model for the corresponding trials and error stripes show the RMSE. For each 5-second time-bin, average performances were compared either between each trial or to zero with paired and single-value Student t-tests. Significance level is provided in the boxes below the plots (ranging from light grey for P<0.05/1 to black for P<0.05/n_Tests_ using Bonferroni’s correction for n_Tests_=3; white for P>0.05). Shaded areas highlight the stimuli critical period based on the on-going distance of the passing virtual shoals illustrated by the fish positions relative to the side screen shown above the time axis.

#### Predator effect

Heatmaps of post-larval fish position density highlight subtle differences in their reaction to dynamic virtual predators versus control parrotfish. With predators, post-larval fish rapidly moved to the upper-right quadrant and remained at the back of the shoal. In contrast, with non-aggressive size-matched controls, post-larval fish also gathered in the upper-right quadrant, but some rapidly started to follow the shoal: group dispersion increased more and earlier than with predators. When both predators and non-aggressive controls were presented (Forced choice), most post-larval fish gathered in the upper-left quadrant with controls, with only a few staying on the predator side or following either predators or controls. The periodic component of the X-position for the Predator trial was smaller and peaked earlier (*A*=36.9 px and *t_0_*=29.7 s) than for Controls (*A*=41.6 px and *t_0_*=32 s, **Table A14**): with bluefin jacks, post-larval fish moved more to the stimulation side and earlier (20-35 s) than with parrotfish (30-35 s, **Figure 5E**). When both fish shoals were presented (Forced choice), X-position of post-larval fish fluctuated around the midline (*b*=–19.4 px and *A*=25.1 px), with a non-significant deviation towards parrotfish. The periodic component of the Y-position peaked for all conditions at similar times (*t_0_*=24.2-29.6 s), with post-larval fish moving rapidly behind fish shoals (**Figure 5F**), especially with predators. However, the combined periodic amplitude and quadratic component in the predator trial, and the linear component in the forced choice task, indicated that, while post-larval fish tended to follow parrotfish and were distributed centrally along the Y-axis at the end of the stimulation (deviating from midline only in the 25-35 s interval), they remained in the upper-half of the aquarium behind virtual predators (Y-position always positive throughout the stimulation). Across the three trials, individual velocities decreased rapidly to about 20 px/s, with troughs at similar times (*t_0_*=22.8-25.8 s). However, model fit of velocity had a quadratic component highlighting that velocity decreased over a longer period of time in the Predator compared to Control trial before increasing again (**Figure 5G; Table A14**). Dispersion decreased earlier than velocities and increased slower with Predators (trough at 35.5 s) than Controls (trough at 31.9 s) although not statistically significant (**Figure 5H**).

## Discussion

Virtual corals – healthy or bleached – displayed on one aquarium side had a significant effect on post-larval fish behaviour (**Figure 3A-D**). Post-larval fish swimming speed increased and the time spent close to the aquarium side displaying the simulated corals increased compared to sand controls with no corals. Interestingly, both healthy and bleached corals attracted post-larval fish. As herbivores, corals are not part of the diet of *A. triostegus*, so this attraction may be due to anfractuosities in the coral framework, potentially providing shelter and/or a hiding place (Leis & McCormick, 2002). In addition, displays of virtual fish highlighted clear and distinct behavioural responses in post-larval *A. triostegus* (**Figure 3E-H**). When presented with five virtual 3D adult conspecifics, post-larval fish were attracted to them within ten to twenty seconds. In contrast, when presented with their natural predators, five virtual bluefin jacks (*C. melampygus*) (Siu et al., 2017), post-larval fish moved to the opposite side of the aquarium with a rapidly decreasing velocity, and then slowly gathered behind the virtual predator shoal. These contrasting responses highlight the ability of these post-larval coral reef fish to visually identify virtual conspecifics and predators and respond differently, with either attraction (conspecifics) or repulsion and/or avoidance (predators), consistent with expectations from natural behaviours. Furthermore, the movement of post-larval fish behind the predator shoal not only confirms their recognition of the virtual predator but also of virtual body features (distinguishing head from tail and positioning themselves accordingly).

The presentation of static or dynamically moving conspecifics led to contrasted reactions in post-larval fish (**Figure 4A-D**). The sudden appearance of a static conspecific shoal startled post-larval fish, causing them to move away or remain stationary, but then they showed attraction, even hitting the side of the aquarium where conspecifics were displayed. However, when the virtual conspecific shoals appeared 2.5 meters away and slowly got closer, no startle responses were observed, rather post-larval fish followed the virtual shoals along the side of the aquarium, even after three identical passes. This dynamic scenario is particularly interesting as it highlights the post-larval fish’s natural cohesive group behaviour, even with virtual conspecifics. Such shoaling and cohesive behaviours with virtual conspecifics have previously been observed in other experiments (e.g., in adult zebrafish *Danio rerio*, Saverino & Gerlai, 2008). In contrast, when static predators suddenly appeared on one aquarium side, post-larval fish slowly moved to the opposite side of the aquarium and gathered behind the virtual predator shoal (**Figure 4E-H**). When virtual bluefin jacks swam by, the reaction of the post-larval fish was less clear across individuals, but mostly consisted of an overall decrease in swimming speed and/or synchronized movements to hide behind the moving virtual predators. When virtual bluefin jacks passed three times, the static-like behaviour became more frequent and individual swimming speeds decreased with each new pass.

We then tested whether these post-larval fish responses were simply due to the size of virtual fish – repulsion to larger fish and attraction to smaller fish – or whether post-larval fish are able to differentiate between virtual fish species. We found subtle yet noticeable differences in post-larval fish reactions to virtual shoals of same-sized non-aggressive controls, butterflyfish *Forcipiger longirostris* and parrotfish *Scarus psittacus*, compared to conspecifics (surgeonfish) and predators (bluefin jacks) respectively. Post-larval fish showed stronger and longer-lasting attraction towards conspecifics with clear shoal-following behaviour and less dispersion, compared to butterflyfish and a preference for conspecifics when both were presented simultaneously (forced choice, **Figure 5A-D**). When presented with predators, the post-larval fish’s back-and-forth escape reaction was triggered earlier, post-larval fish gathered behind the virtual fish for longer and displayed periods with reduced velocity and dispersion at each shoal passage compared to parrotfish (**Figure 5E-H**). In the forced choice task, post-larval fish had a tendency to prefer size-matched parrotfish rather than predators. Altogether, these findings provide evidence that, based on visual cues alone, post-larval fish can distinguish a conspecific from an equally small but non-aggressive fish species, and a predator from an equally large but non-aggressive fish species, suggesting that post-larval surgeonfish respond behaviourally not only to size, but also to the shape and colour pattern of virtual fish.

Over the last two decades, a range of studies have demonstrated that post-larval fish possess developed behavioural and sensory abilities, rejecting the traditional paradigm that they are passive plankton (Leis, 2015; Beldade et al., 2012, 2016). In particular, post-larval fish survival depends on their ability to correctly evaluate sensory cues and select appropriate behavioural responses, *e.g.,* move toward conspecifics or flee predators (Barth et al., 2015). Among the sensory cues used by coral reef post-larval fish, visual cues are the most discussed, but their importance is the least understood (Lecchini et al., 2014). The visual abilities of post-larval fish increase during their pelagic life to reach a maximum near the onset of metamorphosis (Lara, 2001). Our experiments yield convincing and quantified behavioural results highlighting the role of visual cues in post-larval fishes at a stage in their development during which they seek a suitable recruitment habitat. In particular, post-larval *A. triostegus* showed marked attraction towards corals, potentially due to the complex 3D structures with which they are associated. Furthermore, post-larval fish used visual cues to discriminate between conspecifics and predators and tailor their movement and behaviour to either follow their conspecifics (in ecological settings, this could be used to find a suitable settlement habitat) or avoid predation.

## VR for Behavioural Studies

### Proof of concept

The experimental program shows that coral reef post-larval convict surgeonfish (*Acanthurus triostegus*) visually recognize possible hiding places, adult conspecifics, and bluefin jack (*Caranx melampygus*) predators, presented virtually. These results provide a successful proof of concept of our innovative virtual reality setup with automated tracking of fish responses to simulated 3D models of habitats and fish shoals. We overcame two technical challenges: we simulated 3D models of fishes and habitats that were realistic enough to elicit natural reactions in post-larval coral reef fish, and we detected their individual positions in the aquarium in real-time during the trials using a video-based tracking system. The detailed behavioural reactions of *A. triostegus* post-larvae to conspecifics and predators through several relevant kinematic measures such as their position in the aquarium, individual swimming speed, and group dispersion and the multi-factorial analysis of these measures enabled us to disentangle responses that could yield very similar results in less controlled experimental designs (sand *vs*. corals, parrotfish *vs*. bluefin jacks, butterflyfish *vs*. surgeonfish).

### Current limitations and solutions

In this project, we relied on the Microsoft Kinect Azure depth camera, an IR-based technology that proved unsuitable for underwater tracking, forcing us to only use its embedded colour camera. Tracking with a single camera limits the detection of fish in the aquarium to 2D horizontal positions, so we blocked the real-time updating of the viewpoint (see Video-based tracking section). Rendering of the animated scene was however geometrically correct at the centre, and distortions remained limited within the focal aquaria. We believe the behavioural response to the presentation of virtual coral reef habitats would be stronger with the real-time updating of the viewpoint, by enhancing the fish sensation of physically reaching the virtual anfractuosities. To overcome these limitations in our future projects, we recently updated the VR setup to include a pair of high-resolution colour cameras, and developed a complex underwater calibration procedure. Today we can triangulate the detected fish positions from each view, in order to compute in real-time a fish’s 3D position and update the rendering viewpoint in the virtual scene accordingly. Coupling tracking data with the simulation also allows placing the desired test stimulus in the line of sight of focal individuals.

### Perspectives

Numerous other simulations could be used with such experimental setup to test a wide variety of parameters on coral reef fishes at different stages of development, or test coral reef fishes reared in different conditions or exposed to different stresses prior to the visual cue experiment, and quantitatively characterize the impact on their behaviour. The use of VR offers countless new research opportunities including to better understand behaviours of coral reef fish in response to local and global changes (Beldade et al., 2017; Mills et al., 2020; Nedelec et al., 2017; Schligler et al., 2021) and how they impact the role of vision in habitat selection at recruitment. To understand the mechanisms involved in the visual recognition of conspecifics or predators, our VR set-up can be used to manipulate as many visual factors as needed, including size, colour patterns, fin arrangements, but also the behaviour of other individuals (aggressive, curious, social, fleeing). Even if some experimental protocols in aquaria partially control for these factors and facilitate observations compared to *in situ* protocols (e.g. Katzir, 1981; Roux et al., 2016; Besson et al., 2017), the acclimation time required to perform experiments limits the range of possible manipulations. In addition, when real fish are used as stimuli (e.g. Lecchini et al., 2014), the experiment cannot be reproduced multiple times in a reliably comparable manner, as the movement of the stimulus fish cannot be controlled. This study demonstrates that VR, even with a static rendering viewpoint, provides an excellent methodology to apply perfectly-controlled virtual stimuli, which post-larval fish are able to correctly identify. VR can reproduce fairly realistic natural situations that can yield robust statistical results and allow for highly-precise quantifications of post-larval behaviour in response to highly diverse scenarios. In addition, video-based modern tracking technologies have recently emerged in the field of animal behaviour and neuroscience, offering the possibility to conduct fine-grained kinematic analyses of the reactions of animals to specific situations (Portugues & Engert, 2009; Harvey et al., 2009; Barker & Baier, 2015; Dunn et al., 2016; Stowers et al., 2017; Larsch & Baier, 2018; Drew, 2019; Harpaz et al., 2021).

In conclusion, we show proof of concept of our new VR visual stimulation set-up combined with an automated tracking system in an aquarium. The benefits and disadvantages of studying fish behaviour with such technology are listed here. Benefits include shorter habituation phases in the aquarium (approximately 30 seconds, compared to 5-7 minutes when experimenting with non-VR methods, see Nanninga et al., 2017), which we attributed to the non-aggressive immersive baseline environment; a perfect control of visual stimulation (timing and content) allowing for the same stimulations to be repeated within or between individuals several times; the possibility to test the same individuals in multiple successive trials, and in a controlled order; and, as mentioned earlier, the setup enabled us to automatically collect numerous parameters about the kinematics of fish reactions in real-time, which contributed to a precise and objective characterization of their behaviour. Disadvantages include testing the behaviour of fish in a laboratory setting out of their natural milieu, with a visual rendering that cannot reach the quality of a real environment; and the technical skills in computer science required to prepare experiments and to process the large amount of data. However, this successful proof of concept of our new VR setup and automated tracking system on relatively fragile and small post-larval coral reef fish in response to both habitats and different fish species holds significant promise for the field of fish behavioural ecology across all life-stages and fish species in response to multiple biotic and abiotic conditions. The experimental set-up can be scaled up or down as a function of the size of the focal fish and, as such, our VR set-up holds tremendous promise for the future of the study of teleost behaviour.

## Appendices

**Fig A1.**
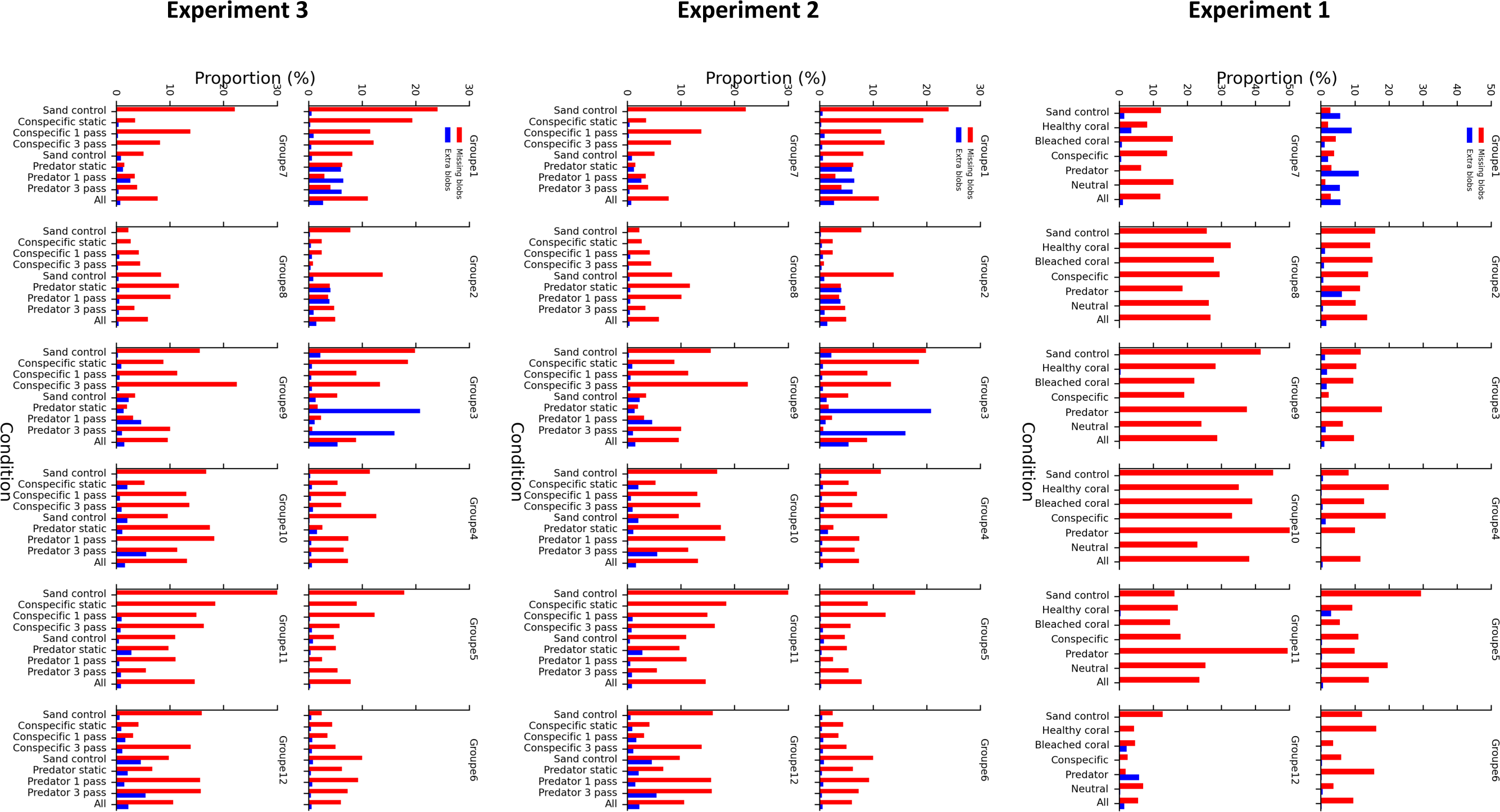
Tracking performance for each experiment and tested fish group in all conditions. The average proportions of missing/extra blobs are given in red/blue. Tracking accuracy was evaluated by computing the average proportion of detected blobs relative to the number of expected blobs (5 per frame) across all recorded frames of the trial. In Experiment 1, on average, 16.3% blobs were missing, and 1.0% extra blobs were detected. The tracking was rather poor for 4 groups (8, 9, 10 and 11), for which there were on average 29.3% missing blobs. However, these groups remained in the analyses as the sampling rate of 5Hz was high enough for the overall data to be valid. In order to improve the tracking performance, we reduced the luminosity variations by using opaque black curtains for the experimental room’s windows. In Experiment 2 and 3, 8.9% and 10.9% of blobs were missing, and 1.5% and 1.3% extra blobs were detected on average respectively.

**Fig A2.**
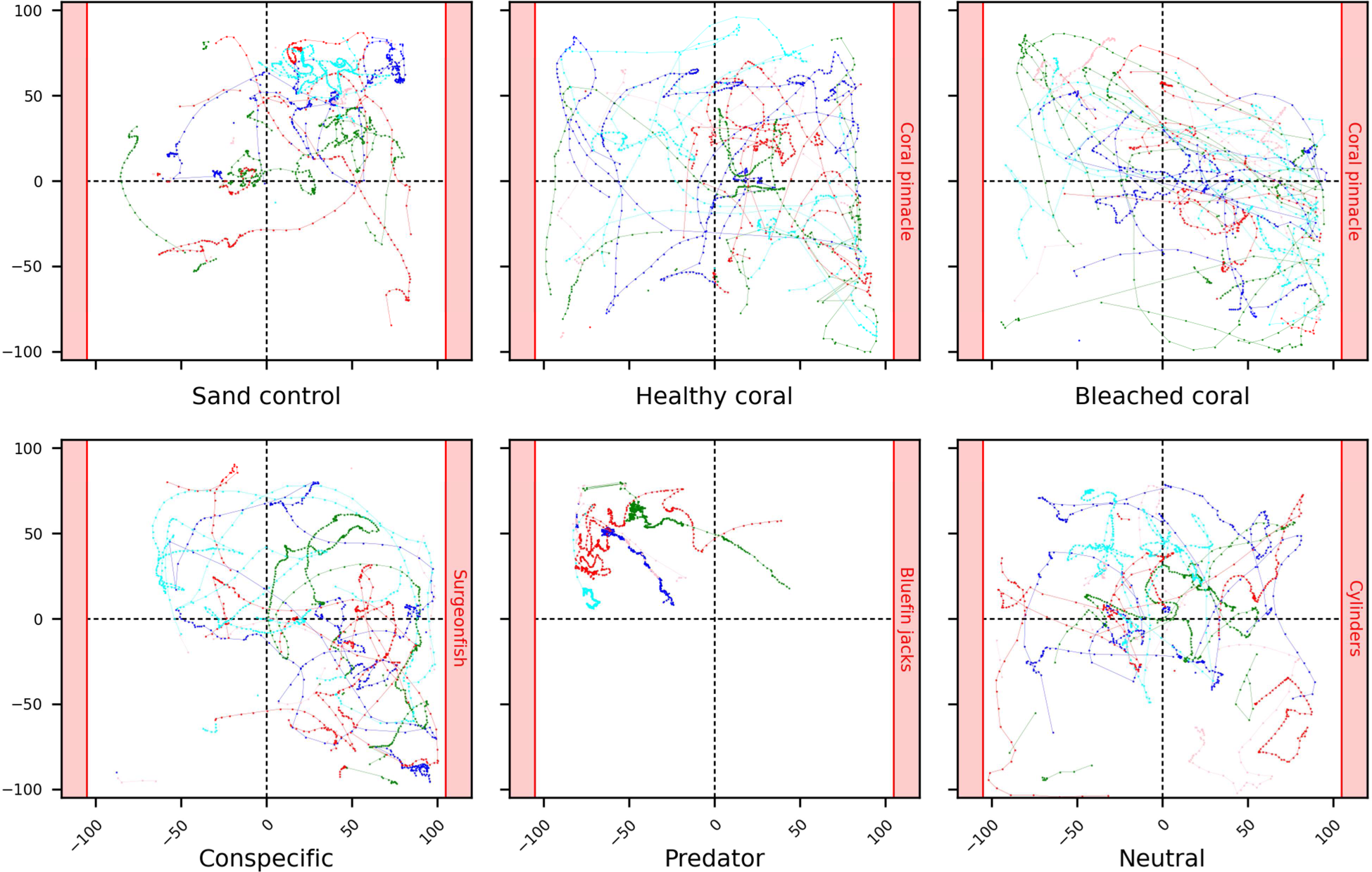
Example of individual trajectories reconstruction in Experiment 1. Individual fish trajectories illustrating behavioural reactions of a typical group to the 6 stimuli presented on the aquarium sides (red shaded areas) of Experiment 1. The validated positions of fish are connected by coloured lines using a minimal distance heuristic to track individual fish in successive frames.

**Fig A3.**
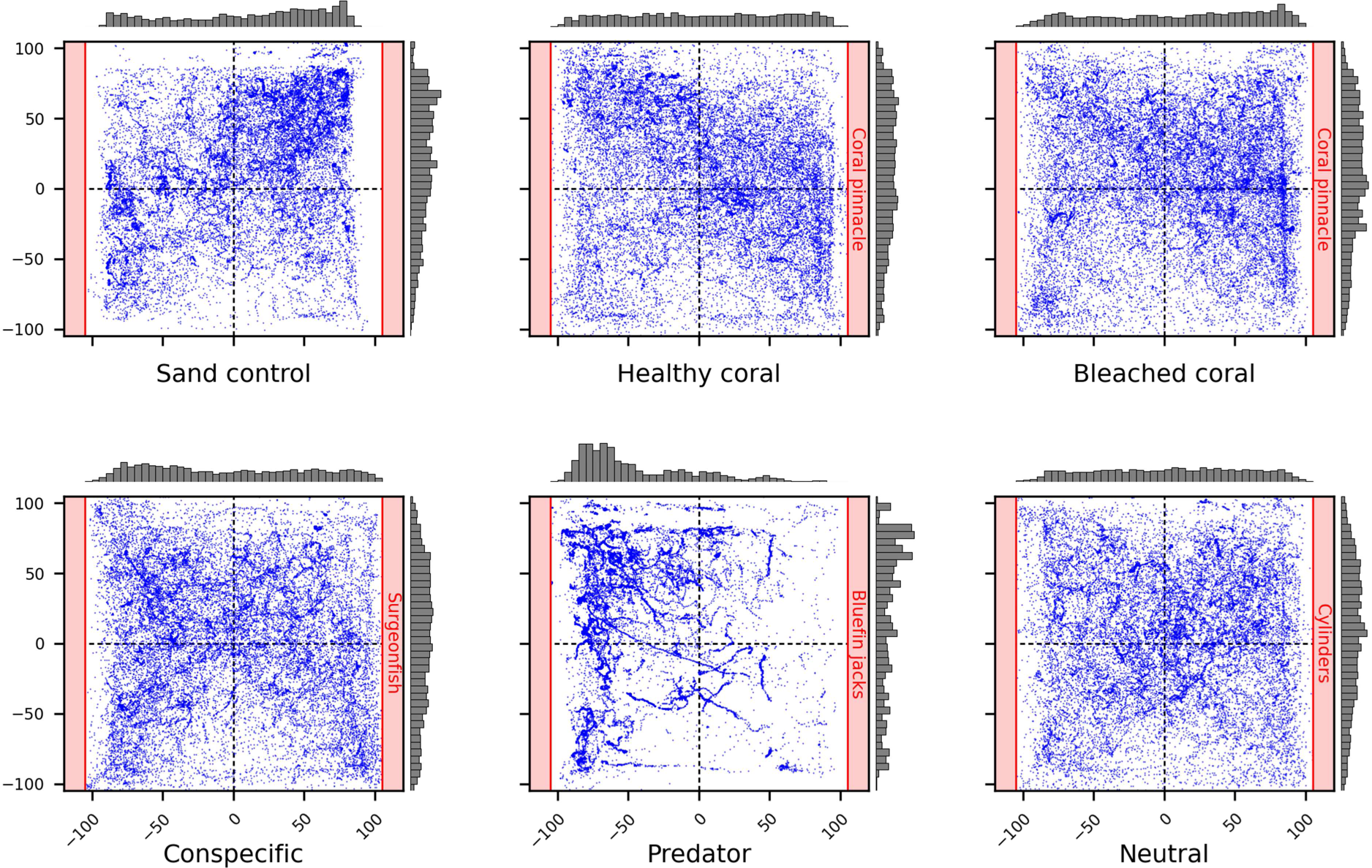
Experiment 1 raw results visualization. 2D positions of all fish and all groups (n=12), in the 6 tested conditions: Sand control, Healthy coral, Bleached coral, Conspecific, Predator, and Neutral. For each condition, the stimulation was presented on the right side. All valid positions detected at each frame throughout trial durations are presented. Normalized X- and Y-position distribution histograms are provided above and to the right of each plot.

**Video A4.**
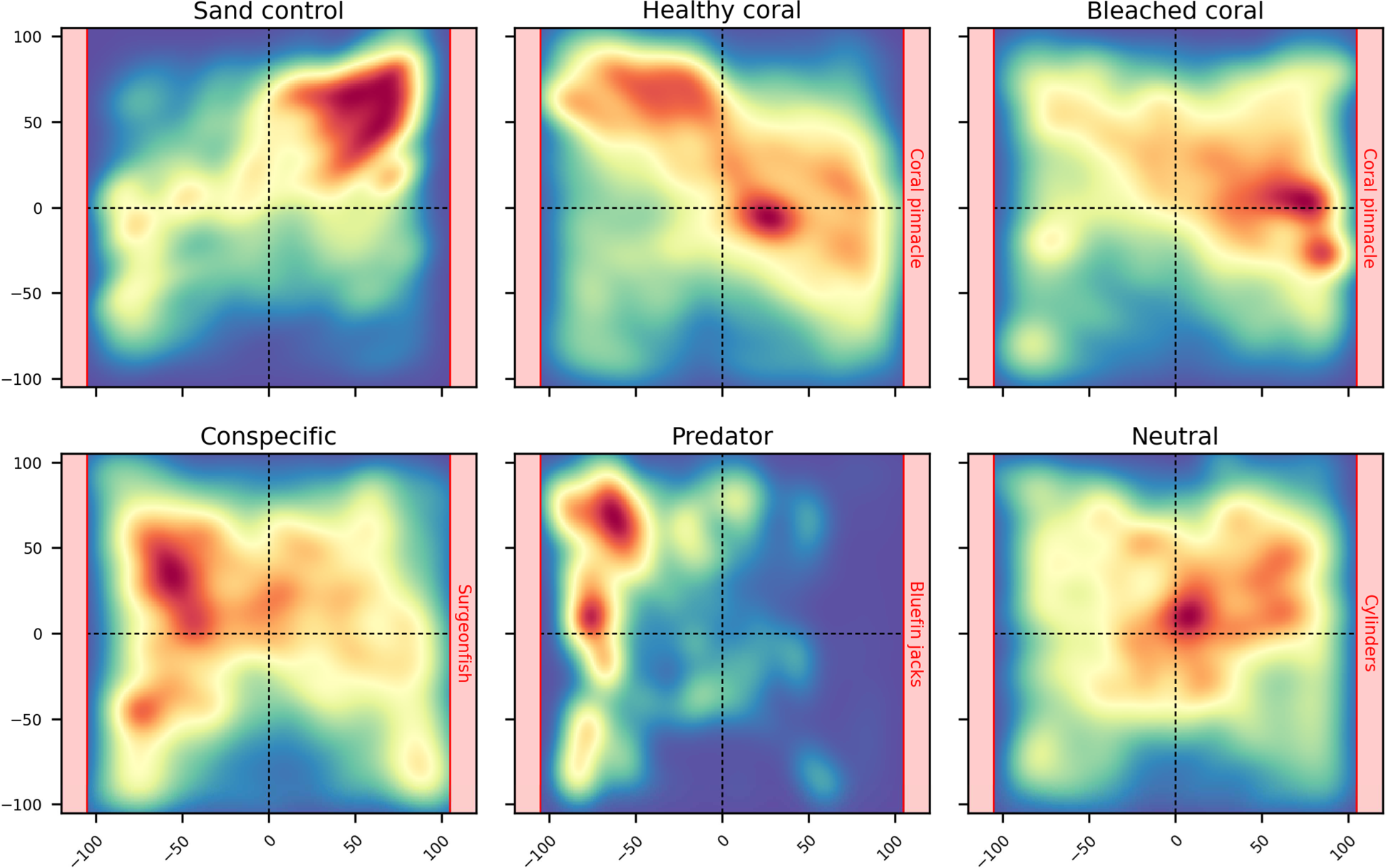
Animation showing the average behavioural responses of Experiment 1. Heatmaps for each condition where fish position density in the aquarium is plotted in successive intervals of 1 second. Density ranges from low (dark blue) to high (dark red).

### Links to videos

×3 (30s): https://amubox.univ-amu.fr/s/C5KfGMSD7mcWfR9

×6 (15s): https://amubox.univ-amu.fr/s/oiaKRwAAqRdeWXw

**Fig A5.**
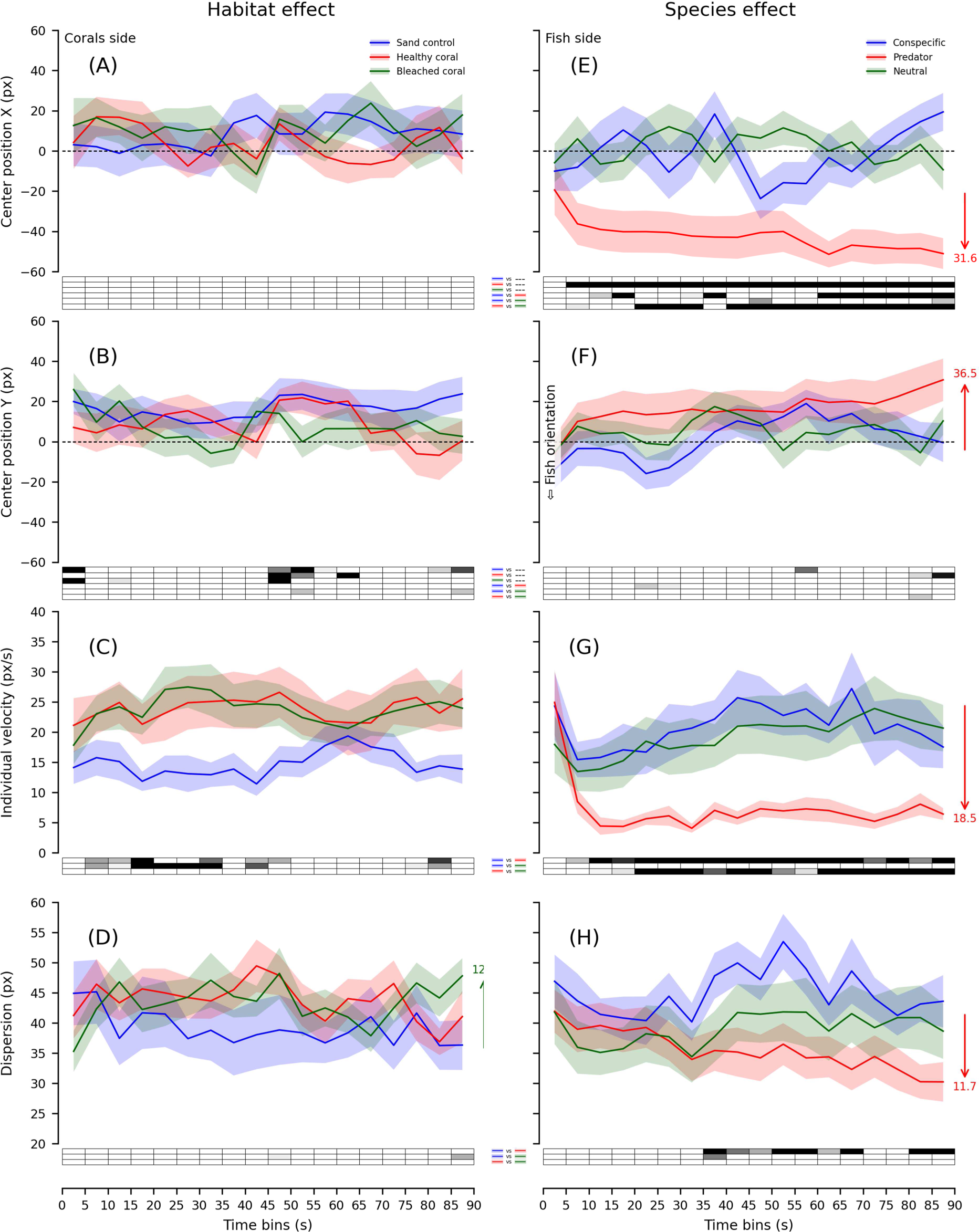
Experiment 1 group time series and statistics. X- and Y-positions of the group centre, individual velocity, and group dispersion plotted in time-bins of 5 seconds for the Habitat effect **(A-D)** and the Species effect **(E-H)**. For each 5-second time-bin, average performance in the conditions was compared to each other or to zero with paired and single-value Student t-tests. Significance level is provided in the boxes below the plots (ranging from light grey for P<0.05/1 to black for P<0.05/n_Tests_ using Bonferroni’s correction for n_Tests_=3; white for P>0.05).

**Table A6.**
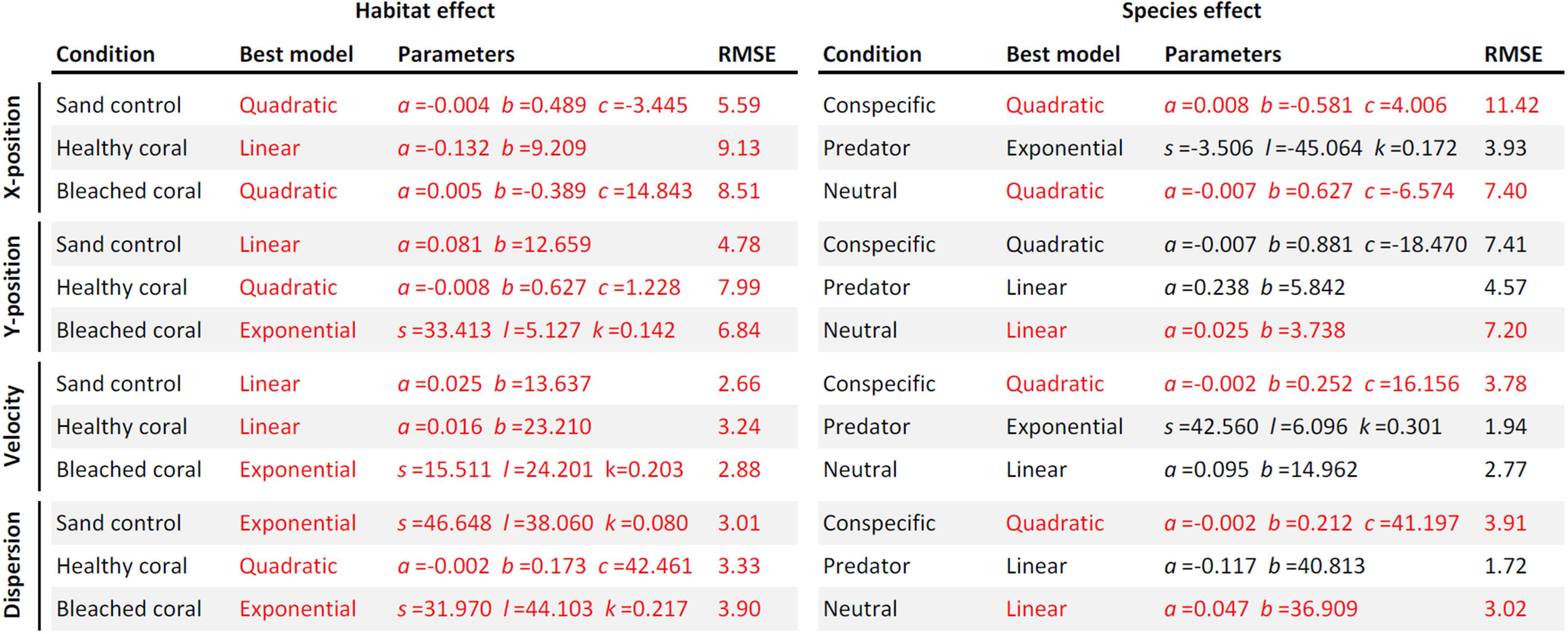
Experiment 1 best behavioural model fitting. The best model, parameters and corrected RMSE for each of the 6 conditions and each of the 4 behavioural measures. For each fit, we assessed the quality of fits computing a RMSE distribution by fitting 1000 times data with shuffled time-bins. Two quality criteria were used: RMSE < dCI(1%) and RMSE+20% < RMSEmean. Fits highlighted in red didn’t meet the quality criteria.

**Fig A7.**
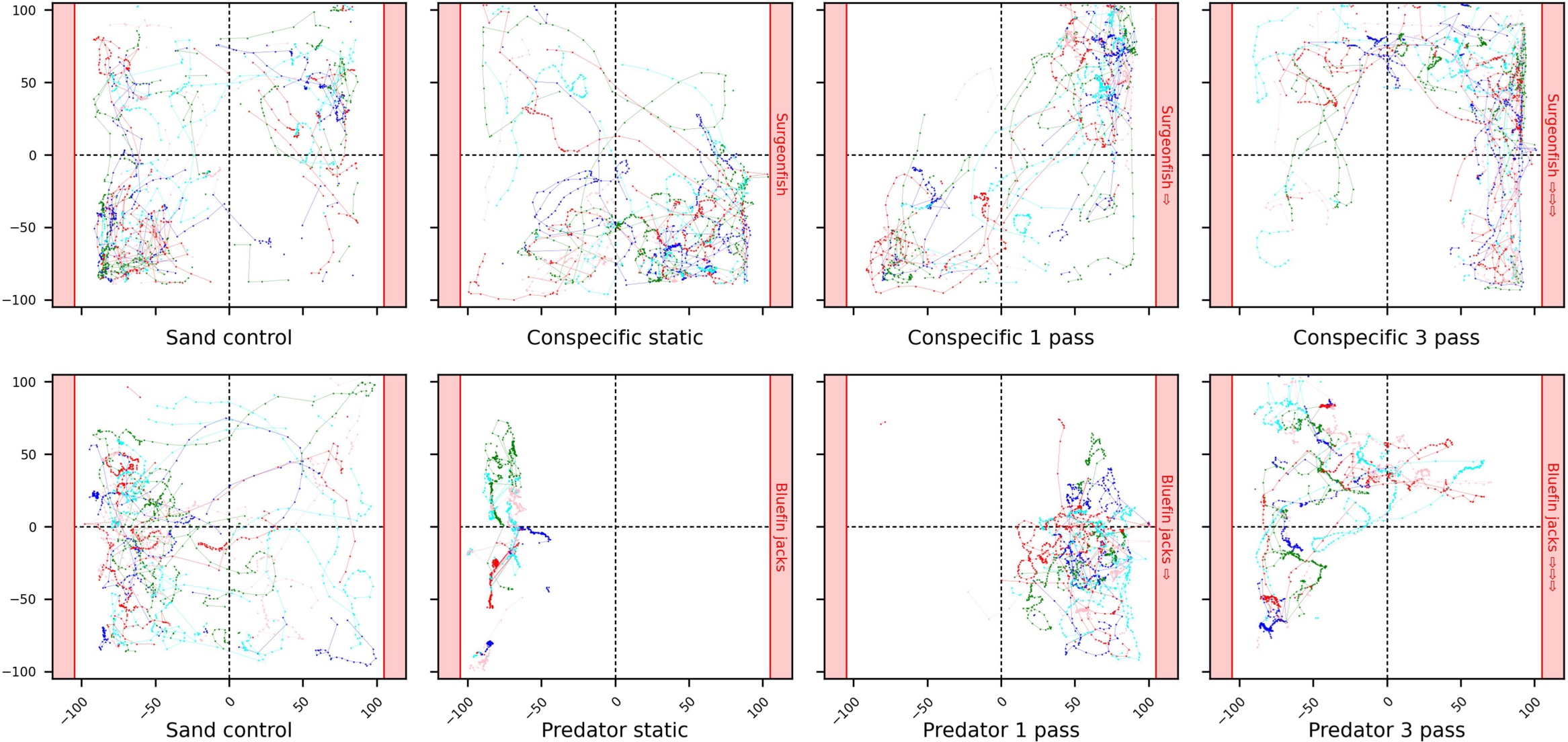
Example of individual trajectories reconstruction in Experiment 2. Individual fish trajectories illustrating behavioural reactions of a typical group to the 8 stimuli presented on the aquarium sides (red shaded areas) of Experiment 2. The validated positions of fish are connected by coloured lines using a minimal distance heuristic to track individual fish in successive frames.

**Video A8.**
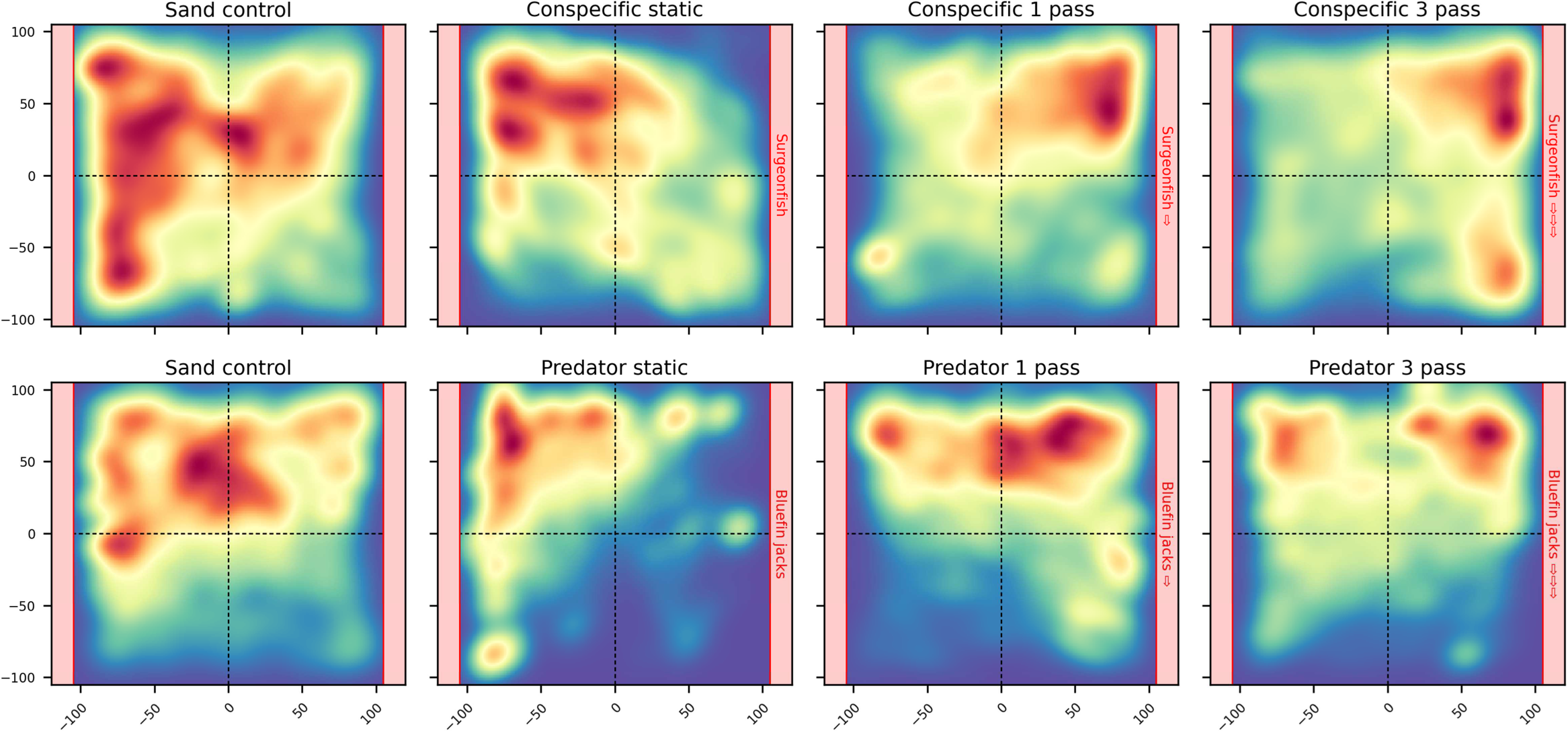
Animation showing the average behavioural responses of Experiment 2. Heatmaps for each condition where fish position density in the aquarium is plotted in successive intervals of 1 second. Density ranges from low (dark blue) to high (dark red).

### Links to videos

×3 (30s): https://amubox.univ-amu.fr/s/F3TWSJoK2PYmP8y

×6 (15s): https://amubox.univ-amu.fr/s/oNojasWfoS6j8K5

**Fig A9.**
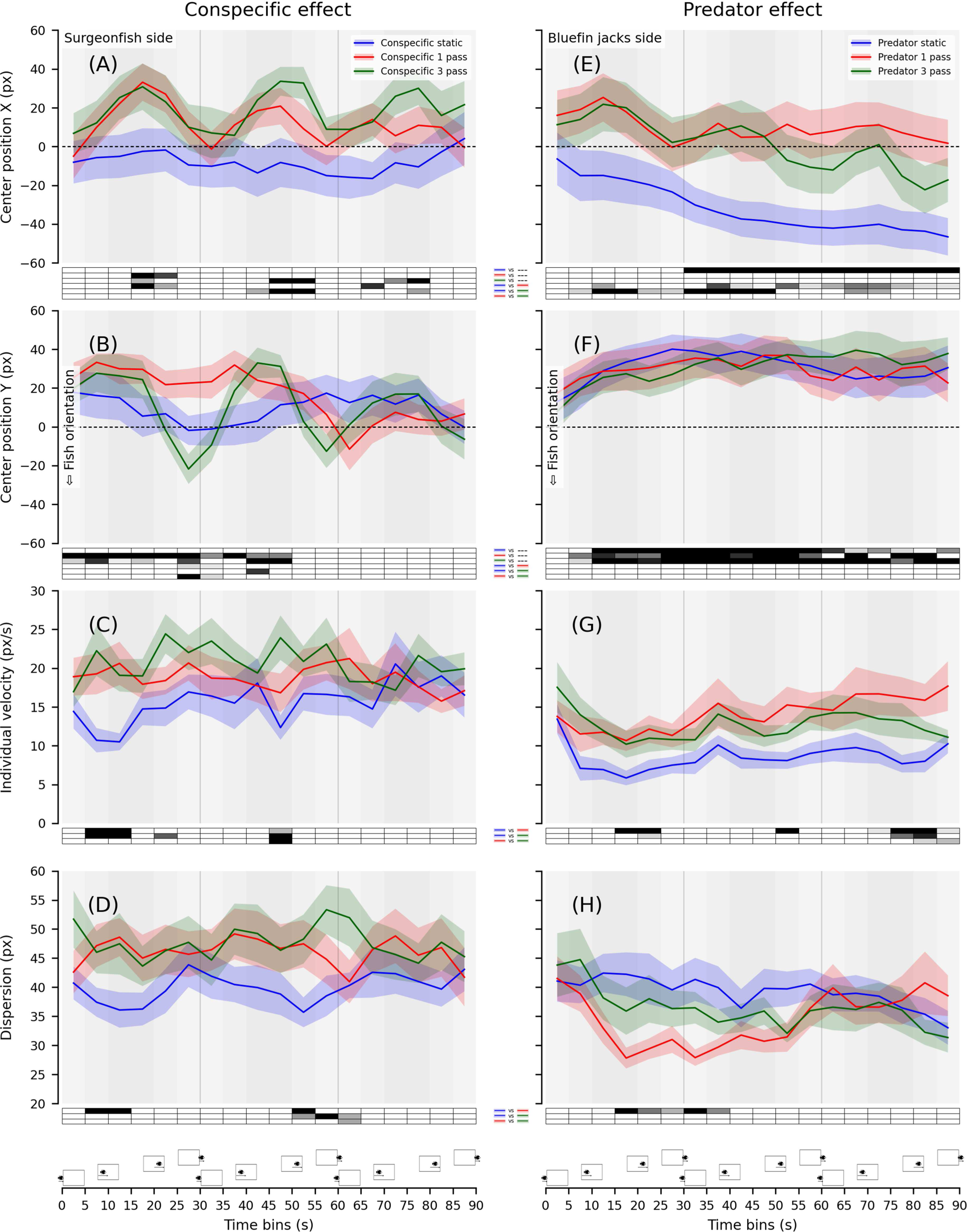
Experiment 2 group time series and statistics. X- and Y-positions of the group centre, individual velocity, and group dispersion plotted in time-bins of 5 seconds for the Conspecific effect **(A-D)** and the Predator effect **(E-H)**. For each 5-second time-bin, average performance in the conditions was compared to each other or to zero with paired and single-value Student t-tests. Significance level is provided in the boxes below the plots (ranging from light grey for P<0.05/1 to black for P<0.05/n_Tests_ using Bonferroni’s correction for n_Tests_=3; white for P>0.05). Shaded areas highlight the stimuli critical periods based on the on-going distance of the passing virtual shoals in the 3-pass conditions (30 s periodicity).

**Table A10.**
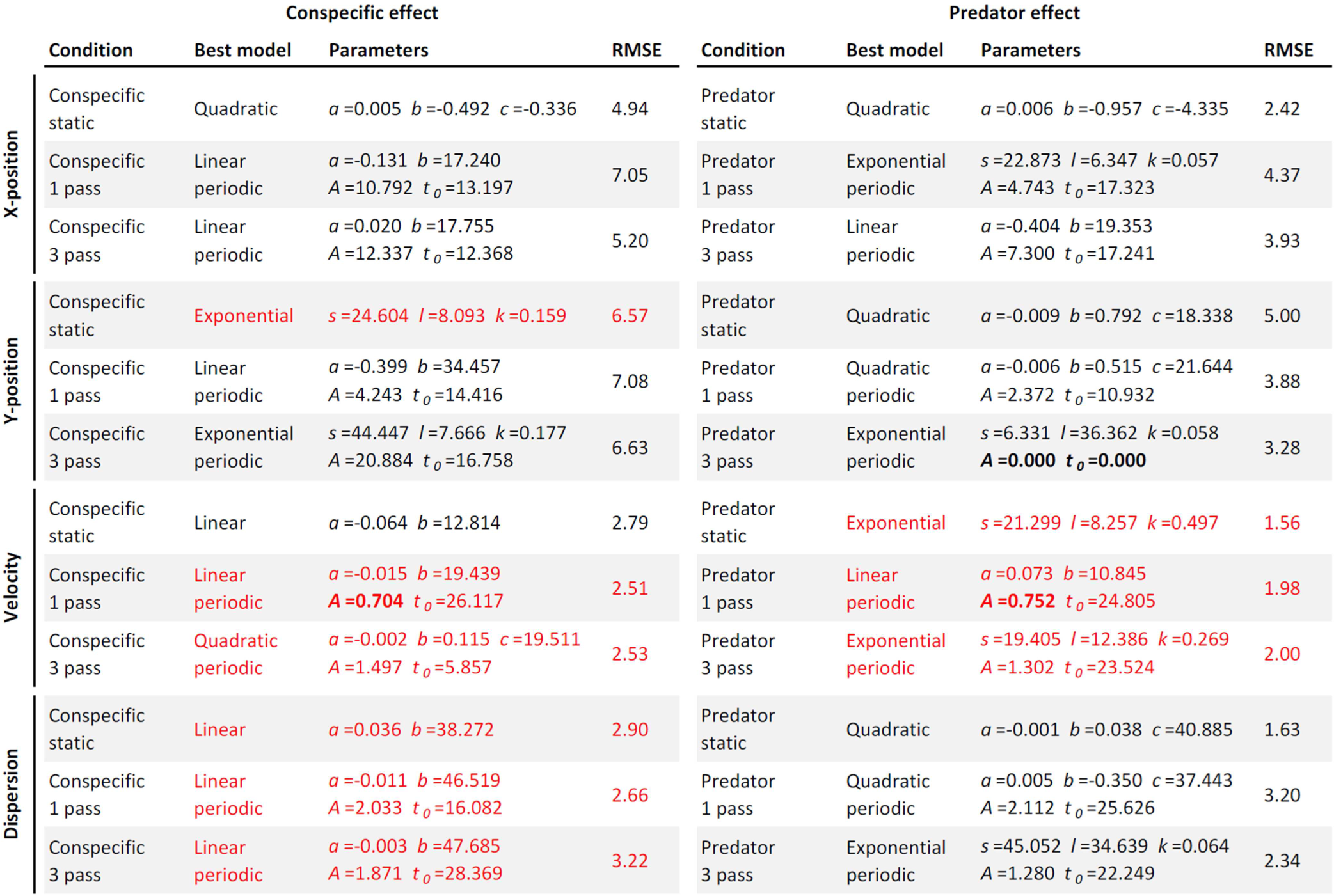
Experiment 2 best behavioural model fitting. The best model, parameters and corrected RMSE for each of the 6 retained conditions and each of the 4 behavioural measures. For each fit, we assessed the quality of fits computing a RMSE distribution by fitting 1000 times data with shuffled time-bins. Two quality criteria were used: RMSE < dCI(1%) and RMSE+20% < RMSEmean. Fits highlighted in red didn’t meet the quality criteria.

**Fig A11.**
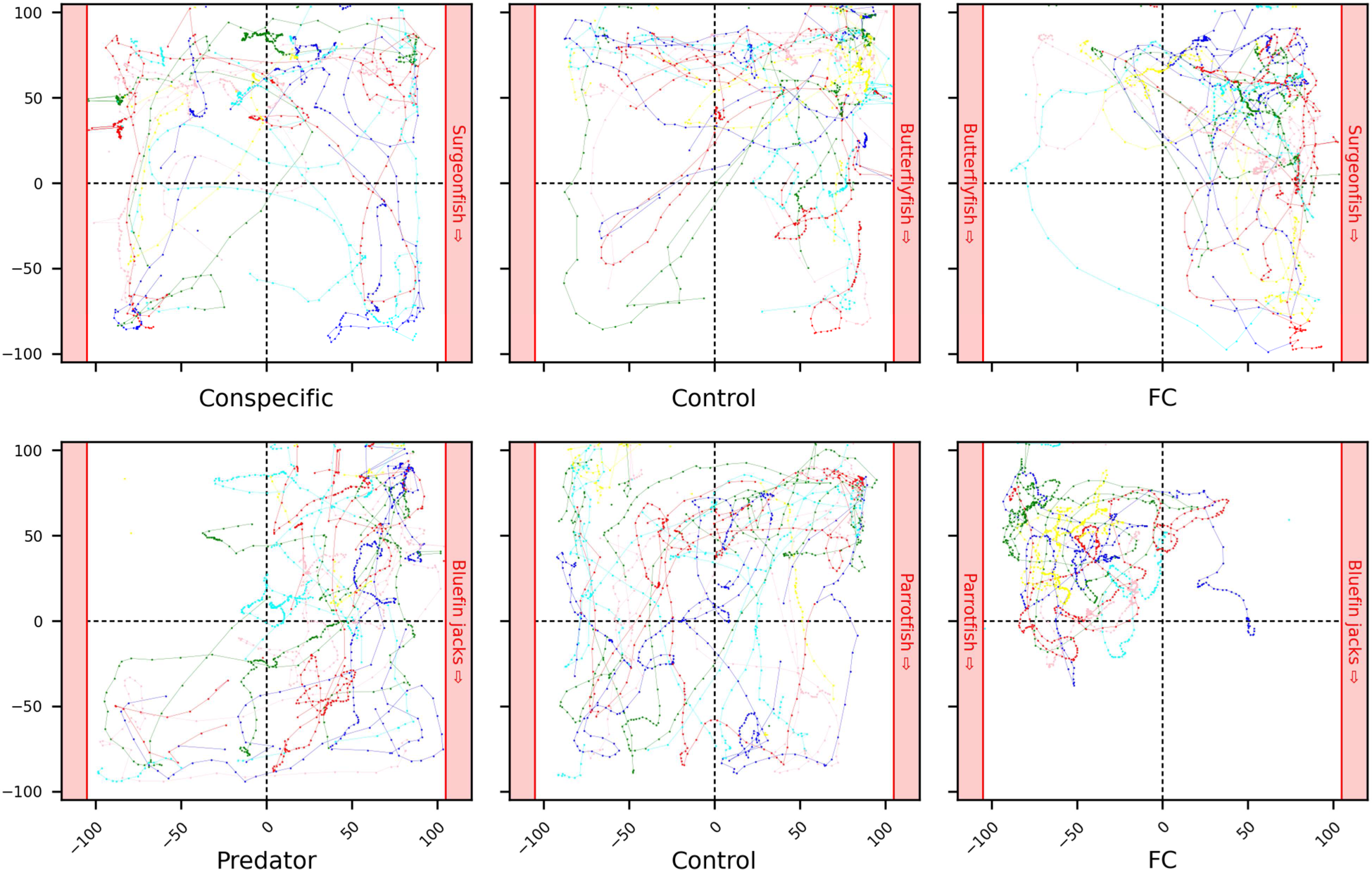
Example of individual trajectories reconstruction in Experiment 3. Individual fish trajectories illustrating behavioural reactions of a typical group to the 6 stimuli presented on the aquarium sides (red shaded areas) of Experiment 3. The validated positions of fish are connected by coloured lines using a minimal distance heuristic to track individual fish in successive frames.

**Video A12.**
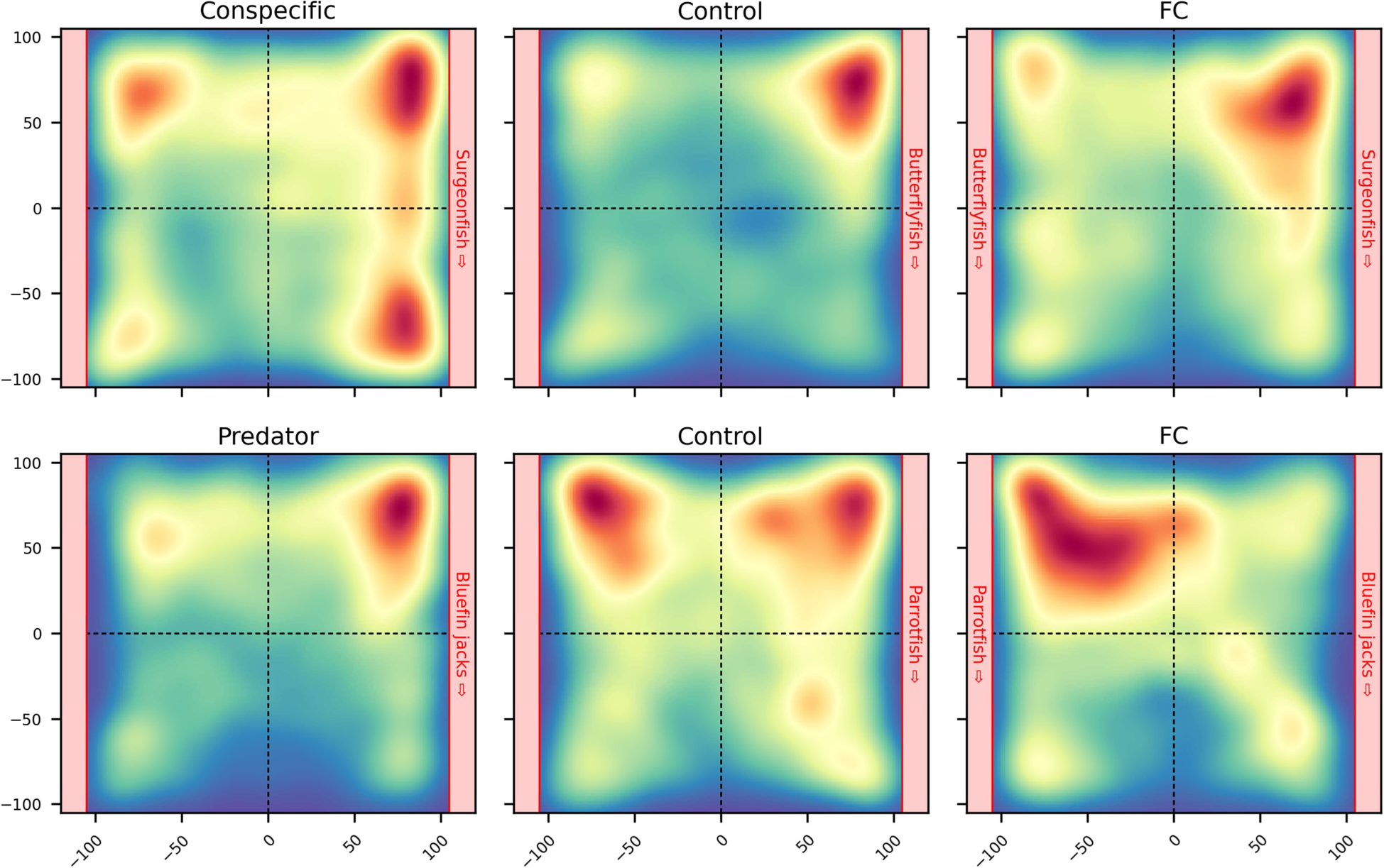
Animation showing the average behavioural responses of Experiment 3. Heatmaps for each condition where fish position density in the aquarium is plotted in successive intervals of 1 second. Density ranges from low (dark blue) to high (dark red).

### Links to videos

×3 (20s): https://amubox.univ-amu.fr/s/32EN2ox6temcaeM

×6 (10s): https://amubox.univ-amu.fr/s/6Zq322jsciYDwka

**Fig A13.**
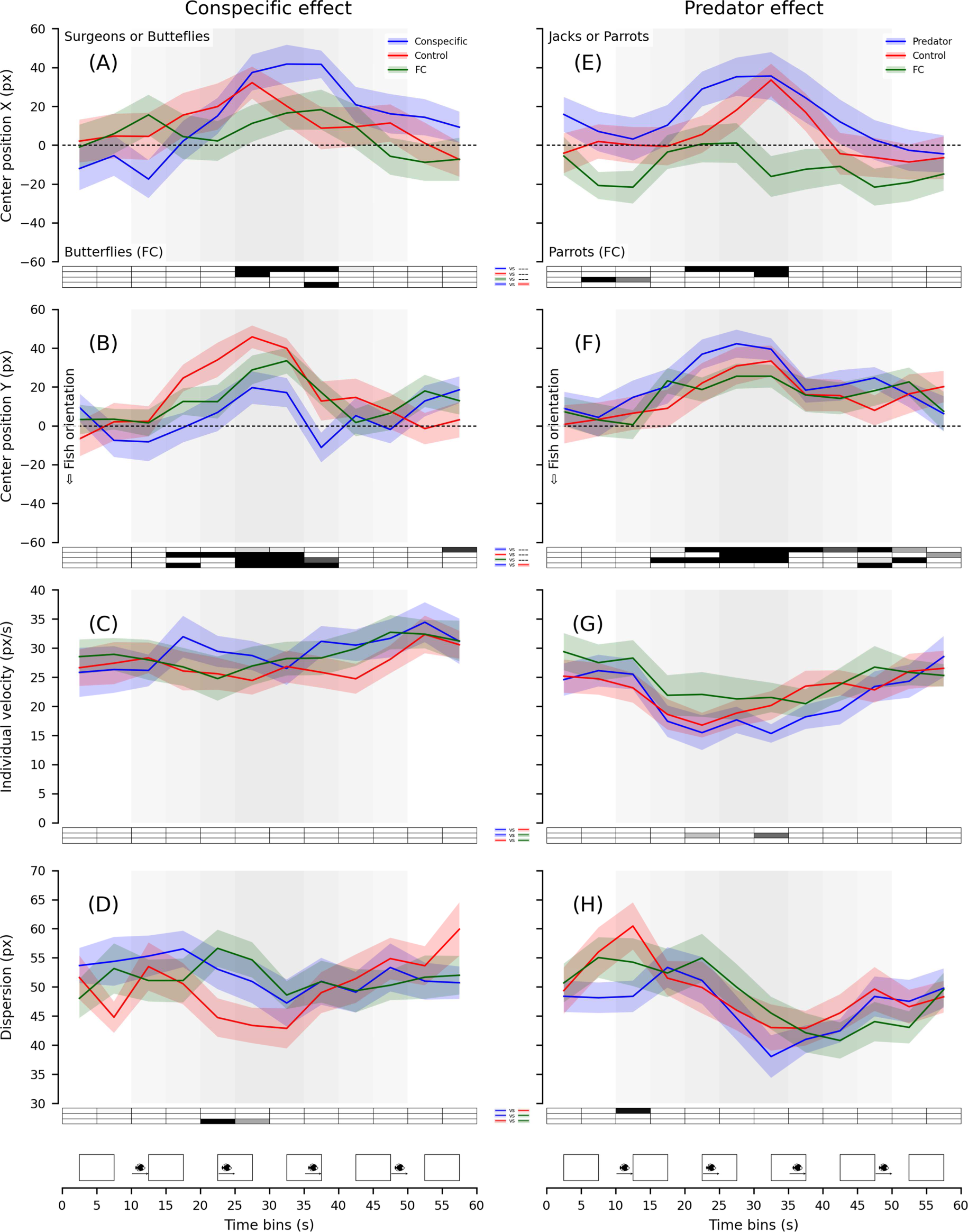
Experiment 3 group time series and statistics. X- and Y-positions of the group centre, individual velocity, and group dispersion plotted in time-bins of 5 seconds for the Conspecific effect **(A-D)** and the Predator effect **(E-H)**. For each 5-second time-bin, average performance in the conditions was compared to each other or to zero with paired and single-value Student t-tests. Significance level is provided in the boxes below the plots (ranging from light grey for P<0.05/1 to black for P<0.05/n_Tests_ using Bonferroni’s correction for n_Tests_=3; white for P>0.05). Shaded areas highlight the stimuli critical period based on the on-going distance of the passing virtual shoals.

**Table A14.**
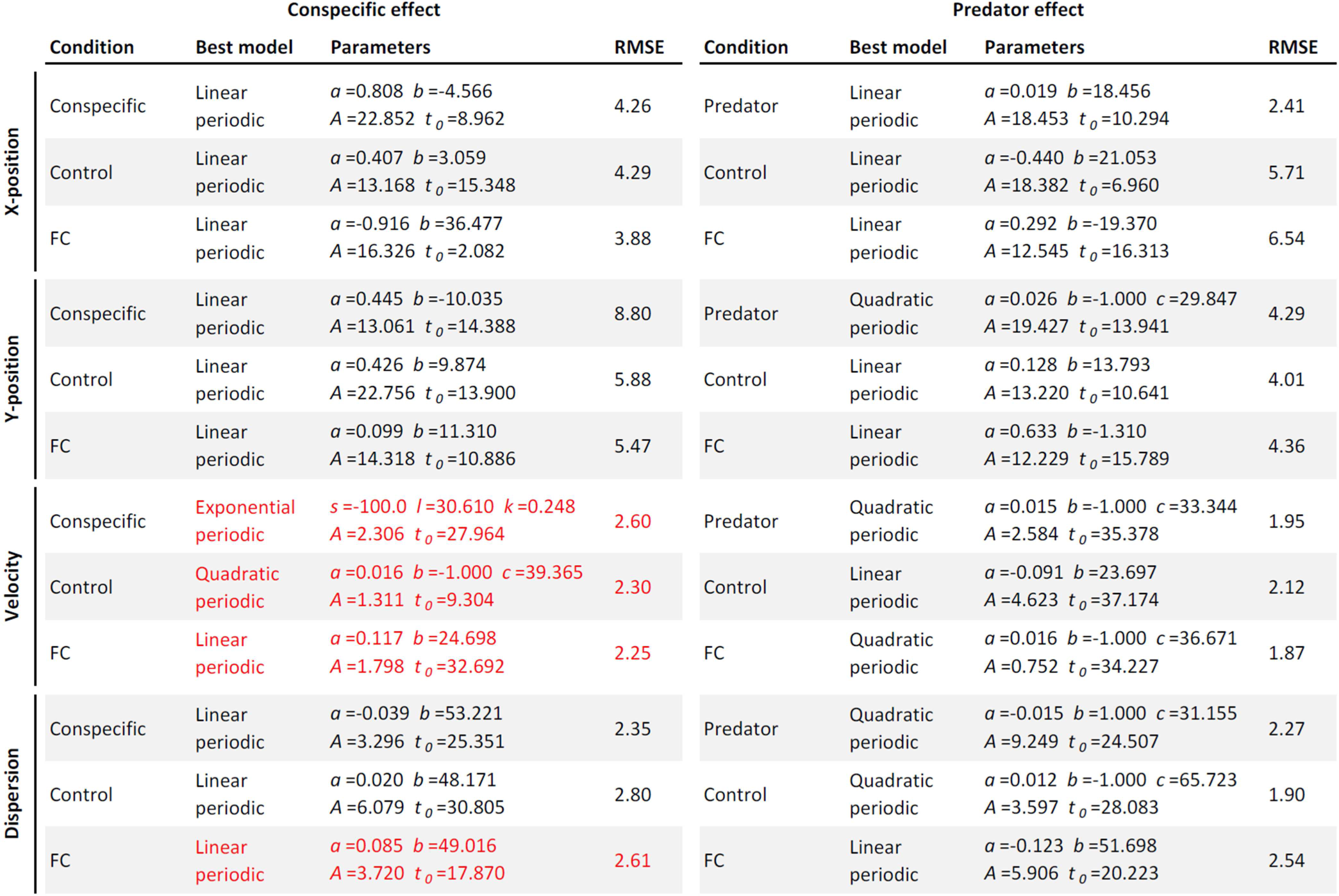
Experiment 3 best behavioural model fitting. The best model, parameters and corrected RMSE for each of the 6 conditions and each of the 4 behavioural measures. For each fit, we assessed the quality of fits computing a RMSE distribution by fitting 1000 times data with shuffled time-bins. Two quality criteria were used: RMSE < dCI(1%) and RMSE+20% < RMSEmean. Fits highlighted in red didn’t meet the quality criteria.

## Data availability

Raw data files, global plots (all groups), and individual plots (trajectories, position scatter plots, animated position density maps, grouped bin data) and the Python code used for data processing for each experiment can be found in the following public repository: https://amubox.univ-amu.fr/s/WyYRBirn3pmzoqT

## Acknowledgements

We would like to thank the technical staff of the CRIOBE for their support, and in particular Matthieu Reynaud for capturing the larvae used in the experiments. This work received support from the French government under the France 2030 investment plan, as part of the Initiative d’Excellence d’Aix-Marseille Université – A*MIDEX, from the Fondation de France (2019-08602), from the Office Français de la Biodiversité (AFB/2019/385 – OFB.20.0888), and from the Agence Nationale de la Recherche (ANR-19-CE34-0006-Manini and ANR-19-CE14-0010-SENSO).

